# Weak local adaptation to drought in seedlings of a widespread conifer

**DOI:** 10.1101/2023.06.09.544307

**Authors:** Rafael Candido-Ribeiro, Sally N. Aitken

## Abstract

There is an urgent need for better understanding how populations of trees will respond to predictable changes in climate and the intensification of extreme weather events such as droughts. The distribution of adaptive traits in seedlings is a crucial component of population adaptive potential and its characterization is important for development of management approaches mitigating the effects of climate change on forests. In this study, we used a large-scale common garden drought experiment to characterize the variation in drought tolerance, growth, and plastic responses to extreme summer drought in seedlings of 73 natural provenances of the two main varieties of Douglas-fir (*Pseudotsuga menziesii* var. *menziesii* and var. *glauca*), sampled across most of their extensive natural ranges. We detected large differences between the two Douglas-fir varieties for all traits assessed, with var. *glauca* showing higher tolerance to drought but slower height growth and less plasticity than var. *menziesii*. Surprisingly, signals of local adaptation to drought within varieties were weak within var. *glauca* and nearly absent within var. *menziesii*. Temperature-related variables were identified as the main climatic drivers of clinal variation in drought tolerance and height growth species-wide, and in height growth within var. *menziesii*. Furthermore, our data indicate that higher plasticity under extreme droughts could be maladaptive in var. *menziesii*. Overall, our study suggests that genetic variation for drought tolerance in seedlings is maintained primarily within rather than among provenances within varieties and does not compromise growth at early stages of plant development. Given these results, assisted gene flow is unlikely to help facilitate adaptation to drought within Douglas-fir varieties, but selective breeding within provenances could accelerate adaptation.

## Introduction

Anthropogenic climate change is transforming and shifting species’ climate spaces at an unprecedented rate, exerting substantial pressure on forests across the globe. With rising temperatures, extreme droughts are growing in frequency, severity, and duration (Dai, 2011; Spinoni et al., 2018; Zhou et al., 2019), and have the potential to disrupt evolved relationships between forest organisms and their environments (Millar & Stephenson, 2015), altering forest functions and composition (Greenwood et al., 2017), and forcing species’ geographical ranges to shift (Parmesan, 2006; Parmesan & Hanley, 2015). Droughts can have detrimental effects on individual trees and on forest stands (Brodribb et al., 2020). From reducing growth rates to driving mortality, there are many direct and indirect mechanisms by which trees can respond to and be affected by drought stress (Hajek et al., 2022; McDowell et al., 2022). Recent evidence suggests, for example, a decline in resilience of temperate forests mainly driven by increasing water deficits and climate variation, overturning possible positive effects on growth from CO_2_ fertilization and warming in past decades (Forzieri et al., 2022). However, predicting stand-level mortality and growth reduction based on knowledge of individual trees’ responses to drought is challenging (Clark et al. 2016; Brodribb et al., 2020). Despite the growing body of studies on the effects of drought on trees and forest dynamics, the study of intraspecific variation in different drought tolerance mechanisms of widespread conifers is still lacking.

Widespread temperate conifers are planted and managed over large geographic areas, and mitigation strategies for the negative impacts of anthropogenic changes could be implemented through planting selected materials or better-adapted ecotypes (i.e., through assisted gene flow (Aitken & Whitlock, 2013)). For that to be effective, a more comprehensive understanding of the climate-driven adaptive responses to climate change in conifers is necessary. Therefore, characterizing the genetic variation of drought tolerance and the potential plastic responses to extreme drought events, as well as testing widespread populations for trade-offs between growth and drought tolerance is crucial to assure that planted stands are going to establish and grow under expected future drought conditions, maintaining ecosystem services.

Divergent selection can generate detectable patterns of local adaptation on adaptive traits, illustrated by the fitness advantage of local populations in their home environment compared to non-local populations (Kawecki & Ebert, 2004). Adaptive differentiation among populations is likely shaped by multiple adaptive traits underlying fitness (Blanquart et al., 2013), and depends on spatially varying environmental conditions and on the interplay between selection and other evolutionary forces, particularly gene flow (Kawecki & Ebert, 2004). Furthermore, local adaptation is an important mechanism by which genetic diversity can be maintained within species (Wadgymar et al., 2022). The study of local adaptation may help reveal the extent of historically maintained variation in increasingly important adaptive traits in tree species, e.g., drought tolerance, and allows the identification of the main selective forces (e.g., climatic variables) driving adaptation. This is particularly relevant in the context of climate change and extreme weather events, as it might unveil populations under higher risk of extirpation and populations with greater potential to tolerate and thrive in future climates. This information can then be used to guide breeding and selection of seed sources for deployment intended to reduce current and predicted future local maladaptation (e.g., Browne et al., 2019).

Local adaptation to climate is common among widespread temperate tree species, despite extensive gene flow (Alberto et al., 2013; Savolainen et al., 2007). More than a century of genecology research on temperate conifers has, for the most part, found significant adaptive clines along provenance temperature-related variables, particularly for growth and cold hardiness (Aitken & Bemmels, 2016; Leites & Benito Garzón, 2023; St Clair et al., 2020). Moreover, cold temperatures were found to be a major limiting factor for growth in temperate and boreal regions (Yang et al. 2023), consistently driving local adaption not only at the phenotypic level but also convergently among species, with distantly related conifer species using some of the same genes to adapt to low temperatures (Yeaman et al., 2016). Precipitation gradients have been found to be weaker drivers of differentiation among populations of temperate conifers (Aitken & Bemmels, 2016; Rehfeldt et al., 2014; St Clair et al., 2005). In spite of that, significant differentiation for drought tolerance has been demonstrated, for example, among small numbers of coastal Douglas-fir (*Pseudotsuga menziesii* var. *menziesii*) populations from contrasting environments (Joly et al., 1989; Kavanagh et al., 1999; White, 1987), suggesting population variation for traits related to drought tolerance.

Studies in recent years have tested larger number of populations of temperate conifers for variation in drought tolerance. Local adaptation to drought has been recently reported in mature trees; for instance, in the easternmost region of white spruce’s range (*Picea glauca*, Depardieu et al., 2020), across most of coastal Douglas-fir’s range (Montwé et al., 2015), and across most of lodgepole pine’s natural range (*Pinus contorta*, Montwé et al., 2016), evidenced by dendrochronological assessments in old common garden experiments. Some of these studies suggested trade-offs between growth and drought tolerance in some regions, indicating that in mature trees, embolism resistance has been selected for over hydraulic conductivity under harsher conditions (Sperry et al., 2008). However, younger trees are in general less resistant to drought than older trees in temperate forests (Au et al., 2022). Moreover, earlier ontogenetic stages (i.e., seedling and sapling) are expected to be the most affected by climate-driven selection pressures (Ramírez-Valiente et al., 2021), allowing populations to respond more quickly to changes in environmental conditions (Williams et al., 2021). Some signals of adaptive differentiation in drought tolerance have also been detected in seedlings, particularly within specific regions of widespread species’ ranges (Bansal et al., 2015a; Csilléry et al., 2020). Nevertheless, range-wide assessments of drought tolerance variation in seedlings of temperate conifers are still largely lacking and are, therefore, urgently needed.

Ideally, local adaptation in trees is assessed via field-based reciprocal transplants of provenances or multiple common garden experiments, from which universal response functions can be estimated (Alberto et al., 2013; Wang et al., 2010). However, these options are constrained by cost and labor requirements to establish, maintain, and assess multiple geographically distant experiments. Furthermore, drought tolerance is difficult to phenotype in large field experiments (Howe et al. 2006). As an alternative, signals of local adaptation can be identified, with relatively high confidence, through the examination of phenotypic clines in adaptive traits along environmental gradients (e.g., climate and geographical), based on a single common garden experiment with seedlings from multiple provenances (Aitken & Bemmels, 2016). Seedlings can, in this way, be experimentally manipulated to allow a direct assessment of the effects of drought. The extent of differentiation for phenotypic traits can be quantified (Q_ST_ or V_POP_) and compared with estimates of differentiation for putatively neutral genetic markers (F_ST_) to assess the strength of local adaptation (Alberto et al., 2013; Whitlock, 2008).

Here, we experimentally investigated the patterns of differentiation in adaptive traits and plastic response to extreme drought conditions in seedlings of the widespread conifer Douglas-fir across most of its natural range and within its two main varieties (*Pseudotsuga menziesii* var. *menziesii* and var. *glauca*). With immensurable ecological significance, Douglas-fir is also one of the most economically important temperate conifer species in the world (Howe et al. 2006; Lavender and Hermann 2014). Naturally occurring from Canada to Mexico and from sea level to over 3,000 m in elevation, Douglas-fir has a natural range spanning approximately 20 million ha in North America (Howe et al. 2006). The large longitudinal, latitudinal, and elevational gradients across its range, as well as the varying topographic features, generate a variety of environmental conditions to which the species has locally adapted over time (Bansal et al., 2015b; Howe et al., 2003; Rehfeldt, 1979; St Clair et al., 2005). This makes Douglas-fir an ideal species for the study of patterns of local adaptation to climate in trees.

In this study, we ask: a) to what extent do Douglas-fir populations from across the species range vary in drought tolerance at the seedling stage, and how strong is the signal of local adaptation to drought species-wide and within-varieties? b) What are the climatic drivers of local adaptation to drought and do they vary between varieties? c) How consistent is the signal of local adaptation across scales of analysis (i.e., species-,and variety-wide)? d) Are there trade-offs between growth and drought tolerance in seedlings? And finally, e) Can plasticity compensate for the lack of drought tolerance?

## Material and methods

### Species, varieties, and sampled provenances

Open-pollinated seedlots were obtained from 73 natural provenances spanning most of the species’ distribution. Seeds were obtained from seedbanks in Canada and USA. We selected from among available seedlots to sample provenances across a wide geographic and environmental range for both the coastal (var. *menziesii*) and the interior variety (var. *glauca*) of Douglas-fir (Table S1; Fig. 1). The sampled seedlots are bulked open-pollinated seeds collected from many trees within each selected provenance for operational reforestation. The seedlots originate from locations spanning a range of more than 16° latitude, 14° longitude and from 61 m to 1737 m in elevation. See Supporting Information for information on how we grew the seedlings prior to the experiment.

**Fig. 1.**
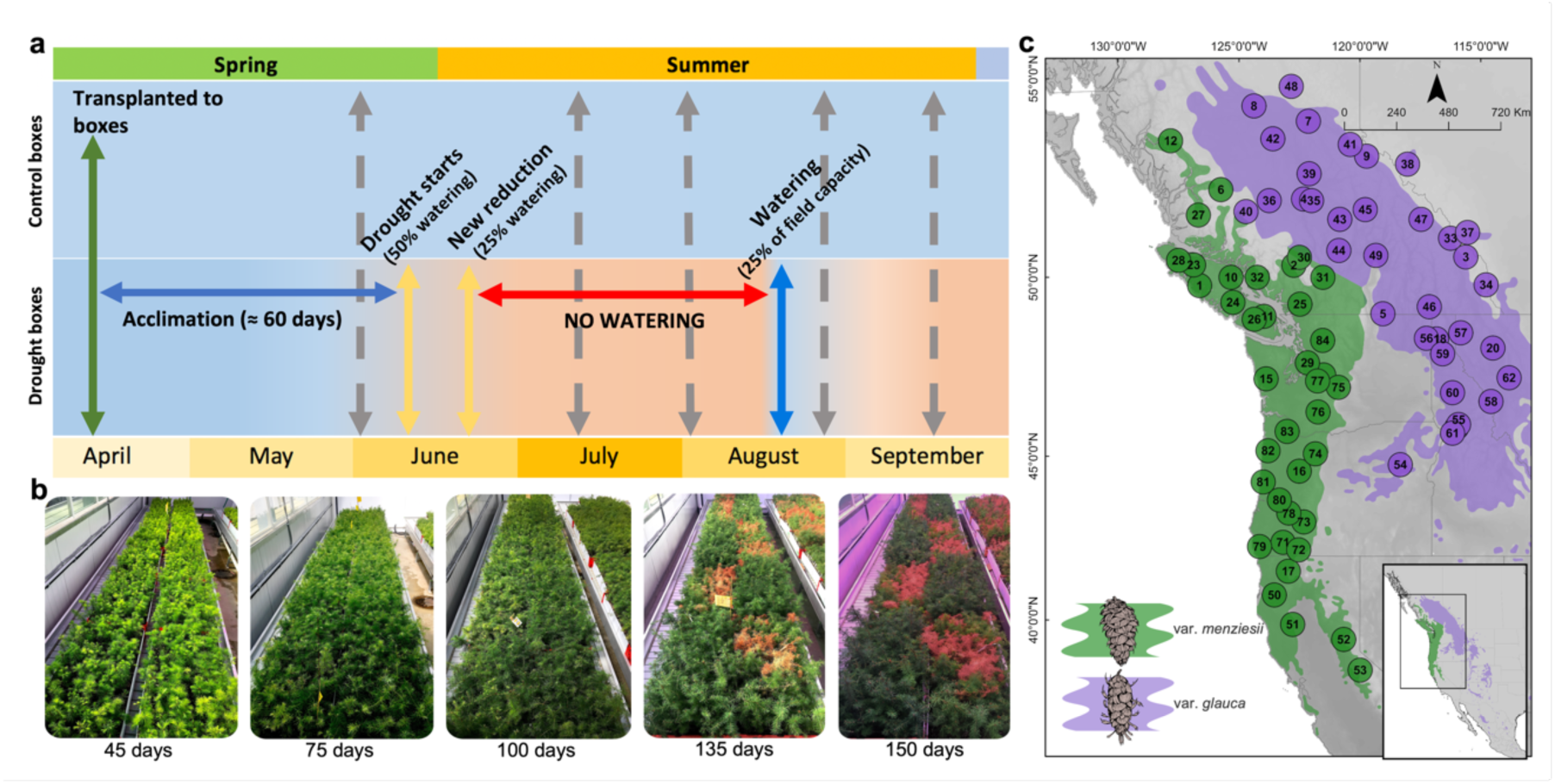
Experimental settings and sampling design. (**a**) Progress of control and drought boxes (treatments) in the experiment over time. During the acclimation phase, all plants were watered as needed. In the drought boxes, yellow arrows indicate partial reductions in the watering regime, the red arrow indicates a complete cessation of watering, and the blue vertical arrow indicates when the drought boxes were re-watered to 25% of field capacity. Grey vertical dashed arrows indicate approximately when all seedlings were assessed for chlorophyll fluorescence (F_v_/F_m_) and height. (**b**) Development of visible drought injury over time, showing strong treatment effects at 135 and 150 days. (**c**) The distribution of the sampled natural provenances across the range of both varieties of Douglas-fir (var. *menziesii* and var. *glauca*).

### Drought to death experiment

The drought experiment was conducted in a greenhouse compartment between April and October 2018 kept at 25° C (± 2° C) during the day and 20° C (± 2° C) overnight, with natural light and photoperiod, in Vancouver, BC, Canada (49.2606° N, 123.2460° W). In April 2018, before bud flush, the seedlings were transplanted into multi-seedling boxes (13 x 40 x 85 cm) made from plastic core board (Fig. 1b) to ensure that all plants within each box experienced similar levels of soil moisture during the drought treatment, regardless of seedling size. This setup also allowed for root competition for water. All boxes were filled with the same volume of commercial substrate (Sunshine^Ò^ Mix #4: organic matter + perlite) and weighed for monitoring and managing subsequent soil volumetric water content. Each box contained 136 seedlings at a 5 x 5 cm spacing. To minimize edge effects, the outer row of seedlings in each box provided a buffer from edge effects and only the 90 central plants were measured.

A split-plot design with single-tree plots and eight slightly unbalanced blocks was used to test all provenances for drought tolerance. Each block was represented by two boxes: one for the drought treatment, and the other as a control treatment, totaling 16 boxes in the experiment. Each provenance was represented by 4 to 19 seedlings in total (average of 16, one seedling per block per treatment), with seedlings randomly allocated to planting locations within each block.

For the first eight weeks after transplanting, we watered the boxes every 5-20 days with tap water and an N-P-K solution (20-8-20 + micronutrients) as needed to maintain soil field capacity. We monitored plant predawn water potential (ll¢_W_) on branches of 3-8 plants per box with a Scholander pressure chamber (3000G4 PWSC) to ensure that ll¢_Wplant_ ≥ -0.5 MPa was maintained. This approach was used to acclimate seedlings and allow root systems to develop. The approximate volumetric water content (VWC) was estimated from box weights every 5-15 days until the end of the experiment. During the acclimation phase, VWC fluctuated between 20% and 45%.

After eight weeks of acclimation, the drought treatment experienced one week of 50% water reduction (relative to the amount to reach field capacity), followed by 75% reduction the following week, and then cessation of all watering. After eight weeks without watering, we provided the seedlings in the drought treatment with a small amount of water once (to 25% of field capacity) to allow surviving seedlings to continue to grow. The same watering regime used during the acclimation period was maintained in the control treatment throughout the experiment to avoid any water stress.

### Measurements

Dark-adapted chlorophyll fluorescence (F_v_/F_m_) was assessed once when the soil was at field moisture capacity, and four additional times over the course of the drought treatment (Fig. 1a), using a chlorophyll fluorometer (FluorPen Z990 – Qubit Systems Inc.). F_v_/F_m_ was used to measure the maximum quantum efficiency of Photosystem II (Murchie & Lawson, 2013) (i.e., photosynthetic efficiency). All the F_v_/F_m_ measurements were performed on needles at night after dark-adapting the seedlings for at least 30 minutes (Murchie & Lawson, 2013). The last F_v_/F_m_ measurement in the control treatment was imputed with the same values from the previous measurement as there was essentially no change in F_v_/F_m_ among the previous measurements (∼ 3% of change on average across individuals), and all measurements were situated within the expected window of minimum variation of F_v_/F_m_ in temperate conifers prior to the transition from summer to fall (Nippert et al., 2004; Yang et al., 2020).

Height increment was used as one measure of drought response. Initial seedling height was measured in the first week after the seedlings were transplanted into the boxes and before bud flush, with five subsequent height measurements during the growing season and experimental drought. The total height increment (THI) for each seedling was calculated as the maximum current year height increment (prior to leader dieback or seedling mortality), and the relative increment was the THI relative to the initial height. The significance of the drought effect on each height increment measurement was tested with the non-parametric Wilcoxon signed-rank test between drought and control treatments implemented in *R 4.2.0* (R Core Team, 2021).

### Geographic and climatic variables

Climatic variables were estimated using the geographic coordinates of each provenance with ClimateWNA (Wang et al., 2016) using the climate normal period of 1961-1990, as this period adequately represents the historic conditions populations are adapted to, and is recent enough that weather stations produced reliable data. The estimated variables were: mean annual temperature (MAT), mean coldest month temperature (MCMT), mean warmest month temperature (MWMT), continentality (TD), extreme minimum temperature over 30 years (EMT), extreme maximum temperature over 30 years (EXT), mean annual precipitation (MAP), mean summer precipitation (MSP), Hargreaves reference evaporation (Eref), and the day of the year on which frost free period ends (eFFP) (Table S1). The same or similar variables were previously found to be relevant for the biology and population differentiation of Douglas-fir (Bansal et al., 2015a; Bansal et al., 2015b; Montwé et al., 2015), and other conifers (Liepe et al. 2016; Yeaman et al. 2016; MacLachlan et al. 2017). The geographic variables latitude, longitude and elevation of provenances were also included in analyses.

## Analysis

### Drought tolerance index (DTI)

To estimate the drought tolerance of each provenance, individual seedling measurements of chlorophyll fluorescence (F_v_/F_m_) were corrected for spatial autocorrelation based on an autoregressive model of residuals implemented in the package *ASReml-R* 4.0 (Butler et al., 2017) (Supporting Information; Fig. S1). A simple linear model was then used to regress the F_v_/F_m_ values of each individual plant corrected for spatial autocorrelation (from here on adjusted values are referred to simply as F_v_/F_m_) on the days of the experiment. Since each plant was modeled individually, we assumed the slope of each linear regression represents the decline in chlorophyll fluorescence of each plant in time, which in turn, reflected the rate of loss of maximum quantum efficiency of Photosystem II due to drought stress (Fig. S2b). The slopes were used in subsequent analyses as the estimated drought tolerance index (DTI). Finally, a mixed-effects model was used for the estimation of best linear unbiased estimates (BLUEs) of DTI for provenances.

To test the significance of treatment effects, we tested the difference between drought and control treatments for individual F_v_/F_m_ measurements, DTI and total height increment using the Wilcoxon rank-sum test, implemented in *R 4.2.0* (R Core Team, 2021). We used this nonparametric test given the different variance structures each treatment produced on phenotypes.

We used the following linear mixed-effects model to test the effect of variety on phenotypes:

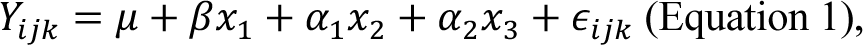

where 𝑌_*ijk*_ is the phenotype corresponding to individuals from variety *i* within one of the treatments (either drought or control treatment), block *j,* and provenance *k*; 𝜇 is the phenotype global mean across all individuals (fixed intercept), 𝛽 is the coefficient for the fixed effect of variety (𝑥_$_), 𝛼_$_ is the random effect of blocks (𝑥_%_), 𝛼_%_ is the random effect of provenance (𝑥_&_) and 𝜖_*ijk*_ is the error term for 𝑌_*ijk*_. Drought treated plants were tested separately from the control plants to avoid violating the assumption of homoscedasticity on linear models.

Differences among provenances within each variety were tested using a simplification of Equation 1, where 𝛽 is the coefficient for the fixed effect of provenance (𝑥_$_) within each variety, 𝛼_$_is the random effect of blocks (𝑥_%_), and 𝜖_*ij*_ is the error term. In this case, the variety term and the random effect of provenances were removed. The significance of the differences observed (fixed effect of varieties or provenance) was tested with Wald Chi-squared tests implemented in the package *ASReml-R* 4.0 (Butler et al., 2017).

### Survival

Using an F_v_/F_m_ of 0.4 as a point of no return, after which plants are unable to recover and die (∼50% reduction in F_v_/F_m_ (Brodribb et al., 2021)), we quantified the proportion of seedlings that had died after every measurement under drought within each variety. Additionally, we used a beta regression modeling approach (Ferrari & Cribari-Neto, 2004), implemented in the R package *betareg* (Cribari-Neto & Zeileis, 2010), to estimate survival of individual seedlings, defined as the approximate day since establishment when each seedling under drought passed the point of no return.

### Phenotypic differentiation (V_POP_)

We partitioned the variance components and estimated standard errors (SE) for each trait by fitting mixed-effects models in *Asreml-R* 4.0 (Butler et al., 2017). The model form used was the same as Equation 1 but with all independent variables set as random effects. We used a similar approach to evaluate the distribution of phenotypic variation within each variety.

We also assessed the differentiation of each phenotypic trait observed between varieties (V_POP-var_) and among provenances (V_POP-prov_) across the species, and among provenances within each variety separately. V_POP_ was calculated by dividing the variance between focal groups (e.g., variety or provenance) by the sum of the variance within (model residuals) plus the variance between the focal groups (adapted from Liepe et al. 2016).

### Phenotypic clines

We used mixed-effects models to obtain the best linear unbiased estimates (BLUEs) of phenotypes (from drought and control treatments separately) for provenances before testing for associations with environmental variables (Supporting Information). Next, using simple linear regressions *R 4.2.0* (R Core Team, 2021), we regressed provenance BLUEs for each trait on each environmental variable for the two varieties combined and individually. We tested both linear and quadratic models and used a false-discovery-rate-adjusted *p-value* (*q-value*) to account for multiple comparisons.

### Plasticity and trade-offs

The pattern and degree of plasticity for height growth in response to drought were examined based on reaction norms constructed for each provenance based on seedlings growing in control and drought treatments (Pigliucci 2001). The degree of plasticity of each provenance and variety was defined as the difference between the relative height increment BLUE estimated for each of these groups in the control treatment and in the drought treatment. The pattern of plasticity for each provenance was defined as the sign of the difference (Pigliucci, 2001). The significance of the differences was tested with non-parametric Wilcoxon Test.

To examine possible trade-offs between drought tolerance and height growth in Douglas-fir seedlings, which in turn could reflect a hydraulic safety-hydraulic efficiency trade-off, simple linear regressions were performed at the species-wide and within-variety levels, with total height increment BLUEs as the dependent variable and drought tolerance index (DTI) BLUEs as the independent variable. We also used simple linear regressions to test for plasticity-survival trade-offs at the species-wide and within-variety levels, with degree of plasticity as the dependent variable and survival BLUEs as the independent variable.

## Results

### Effects of drought on photosynthetic efficiency

For the first measurement (53 days after establishment and 16 days before drought started), F_v_/F_m_ did not differ between the two treatments at the species-wide level nor within varieties (Table S2). Seedlings in the drought treatment started diverging significantly from those in the control from the second measurement (29 days of drought) onwards, as water deficits increased in the drought treatment. All subsequent individual measurements showed significant effects of drought compared to controls, with a significant reduction in the photosynthetic efficiency both species-wide and within varieties (Table S2).

Variety *glauca* showed consistently higher F_v_/F_m_ than var. *menziesii* for all measurements in both drought and control treatments (Fig. 2c; Fig S8; Table S3). The population differentiation observed between varieties (V_POP-var_) across all individual F_v_/F_m_ measurements in the drought treatment varied between 0.14 and 0.45, with the second measurement (29 days of drought) showing the greatest differentiation between varieties (Table 1). The substantial variation in F_v_/F_m_ explained by provenances species-wide during the second and third F_v_/F_m_ measurements in the drought treatment (V_POP_=0.32 and 0.27 respectively) far exceeds the genome-wide population genetic structure estimated for the same provenances (F_ST_ = 0.107, Table 1).

**Fig. 2.**
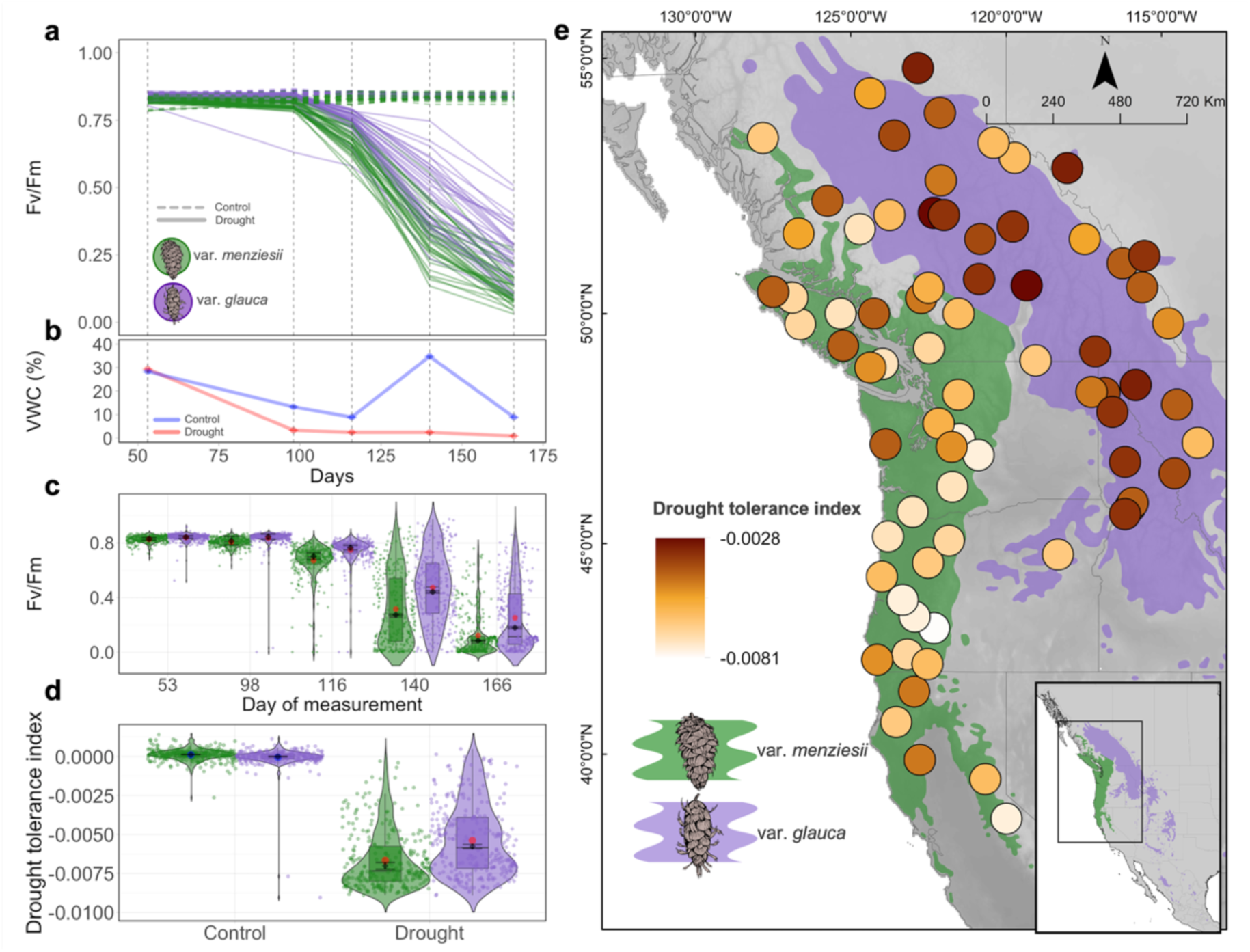
Drought tolerance in seedlings of 73 natural provenances of Douglas-fir. (**a**) Average provenance trajectories of photosynthetic efficiency (F_v_/F_m_, corrected for spatial autocorrelation) through time in control (horizontal dashed lines) and drought (solid lines) treatments. (**b**) Average soil volumetric water content (VWC (%)) among control (blue) and drought (red) treatments. Vertical grey dashed lines in **a** and **b** indicate timing of measurements. (**c**) The five individual F_v_/F_m_ measurements of all var. *menziesii* (green) and var. *glauca* (purple) seedlings (points) after correction for spatial autocorrelation. (**d**) Drought tolerance index (DTI; estimated slopes of F_v_/F_m_ regressed on day of experiment) in control and drought treatments. Red (in **c** and **d**) and blue (in **d**) points depict the absolute trait averages of each variety among drought and control treatments, respectively. Diamonds depict the best linear unbiased estimated values (BLUEs) for each variety and treatment, and error bars are the standard errors of the BLUEs. (**e**) Distribution of DTI across all provenances. More negative values in **d** and **e** indicate less drought tolerance.

**Table 1.**
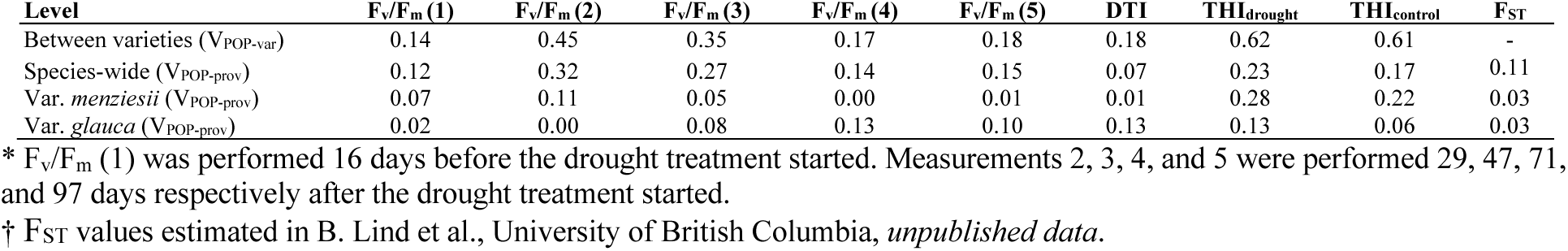
Differentiation between varieties (V_POP-var_), and among provenances (V_POP-prov_) within species and varieties for all F_v_/F_m_ measurements (1-5)* after correction for spatial autocorrelation in the experiment, drought tolerance index (DTI, estimated as the slopes of the F_v_/F_m_ values regressed on the days of the experiment) in the drought treatment, and total height increment (THI) in drought and control treatments. F_ST_, estimated genome-wide genetic structure^†^.

Within var. *glauca*, significant differences were detected among provenances in the drought treatment from the third measurement on (Table S4). The largest population differentiation in F_v_/F_m_ among var. *glauca* provenances (V_POP-prov_) was for the fourth measurement, 71 days after the drought commenced (V_POP-prov_ = 0.13) (Table 1). In contrast, within var. *menziesii*, differences were detected in the first measurement (before the drought treatment) and persisted only until the third measurement under drought (Table S4). The largest F_v_/F_m_ differentiation among var. *menziesii* populations was produced by the second measurement (V_POP-prov_ = 0.11) (Table 1).

A highly significant effect of drought was also observed for the drought tolerance index (DTI), estimated as the slope in decline in F_v_/F_m_ after correction for spatial autocorrelation, indicating an overall rate of reduction in photosynthetic potential due to drought stress (Fig. 2d). At the species-wide level, the estimated DTI for the drought treatment was significantly lower (−0.005971) than the near-nil value for the control (0.00004) (*p* <<0.0001; Fig. S6). Variety *glauca* had a significantly slower decline in F_v_/F_m_ DTI under drought (DTI = -0.00578, *p* <<0.0001) than var. *menziesii* (DTI = -0.007019; Fig. 2d).

Surprisingly, no significant differences in DTI were observed among var. *menziesii* provenances (*p* = 0.449; Table S4), and the V_POP-prov_ observed within this variety was negligible (Table 1). Meanwhile, a highly significant difference among var. *glauca* provenances was observed for the same trait (*p* <0.0001; Table S4), with provenances varying from -0.007428 to -0.002819. A V_POP-prov_ of 0.13 was observed within var. *glauca* (Table1).

### Survival

The estimated proportion of dead individuals (F_v_/F_m_ < 0.4) was negligible in the first two measurements, but varieties started to differentiate after the third measurement (47 days after drought), with 6% of var. *menziesii* and 2.5% of var. *glauca* individuals dying (Fig. S12). After 71 days of drought, var. *menziesii* had 64% mortality and var. *glauca* had 41%. In the last measurement, 97 days after the drought treatment started, 92% of var. *menziesii*, and 74% of var. *glauca* individuals were dead.

### Growth responses to drought

The drought treatment reduced seedling growth. The estimated total height increment (THI) in the drought treatment was significantly lower (7.00 cm, *p* << 0.0001) than in the control (8.32 cm; Table S2). Within both varieties, the difference in THI between control and drought treatments was also significant (var. *menziesii p* <<0.0001; var. *glauca p* = 0.00127; Table S2). The two varieties also differed, with var. *glauca* plants growing less on average in both treatments (THI_dry_ = 5.22 cm; THI_wet_ = 5.74cm) than var. *menziesii* (THI_dry_ = 9.08 cm; THI_wet_ = 11.35 cm; *p_dry_* and *p_wet_* <<0.0001; Table S3). Even though the effect of drought was greater on coastal provenances, with a 20% reduction in growth, they still produced, on average, taller seedlings than the interior provenances, which had a 9% reduction in growth under drought. Moderate differentiation (V_POP-prov_) for THI in drought (0.28) and control (0.22) treatments were observed among var. *menziesii* provenances. In contrast, within var. *glauca*, V_POP-prov_ for THI in drought and control treatments were much lower (0.13 and 0.06, respectively; Table 1).

To characterize the effects of drought on the timing and duration of growth, height was measured five times during the experiment in both treatments. Overall, height increment in the drought treatment was significantly lower than in the control for the second and third measurements, 29 and 47 days after the drought started (*p* = 0.00019, and *p* <<0.0001, respectively). All seedlings in both treatments had ceased growth by subsequent measurement dates (71 and 97 days after the drought started), so there were no differences between treatments in growth during this period (Table S2). The reduction in growth was first observed for var. *menziesii*, 29 days after the drought treatment started (*p* <<0.0001). No significant effect of drought on height increment was observed within var. *glauca* until the third measurement, 47 days after the start of the drought treatment (*p* = 0.0012; Table S2).

### Phenotypic clines along climate gradients

Consistent with the weak signal of local adaptation to drought, linear relationships of drought tolerance index (DTI) with ten provenance climatic and three geographic variables were also weak but in general significant (Fig. 3; Table S5). DTI was most strongly associated with the provenance climatic variables continentality (TD; R^2^ = 0.36, *q* < 0.0001), mean coldest month temperature (MCMT; R^2^ = 0.26, *q* < 0.0001), and extreme minimum temperature over 30 years (EMT; R^2^ = 0.22, *q* = 0.0001). Surprisingly, the weakest significant relationship observed was with Hargreaves reference evaporation (Eref, R^2^ = 0.08, *q* = 0.0221), and no significant association of DTI was detected with mean summer precipitation (MSP; R^2^ = 0.00, *q* > 0.99). DTI was positively associated with provenance latitude, longitude, and TD, and negatively associated with MCMT, EMT, mean annual temperature (MAT), end of frost-free period (eFFP), mean annual precipitation (MAP), and Eref.

**Fig. 3.**
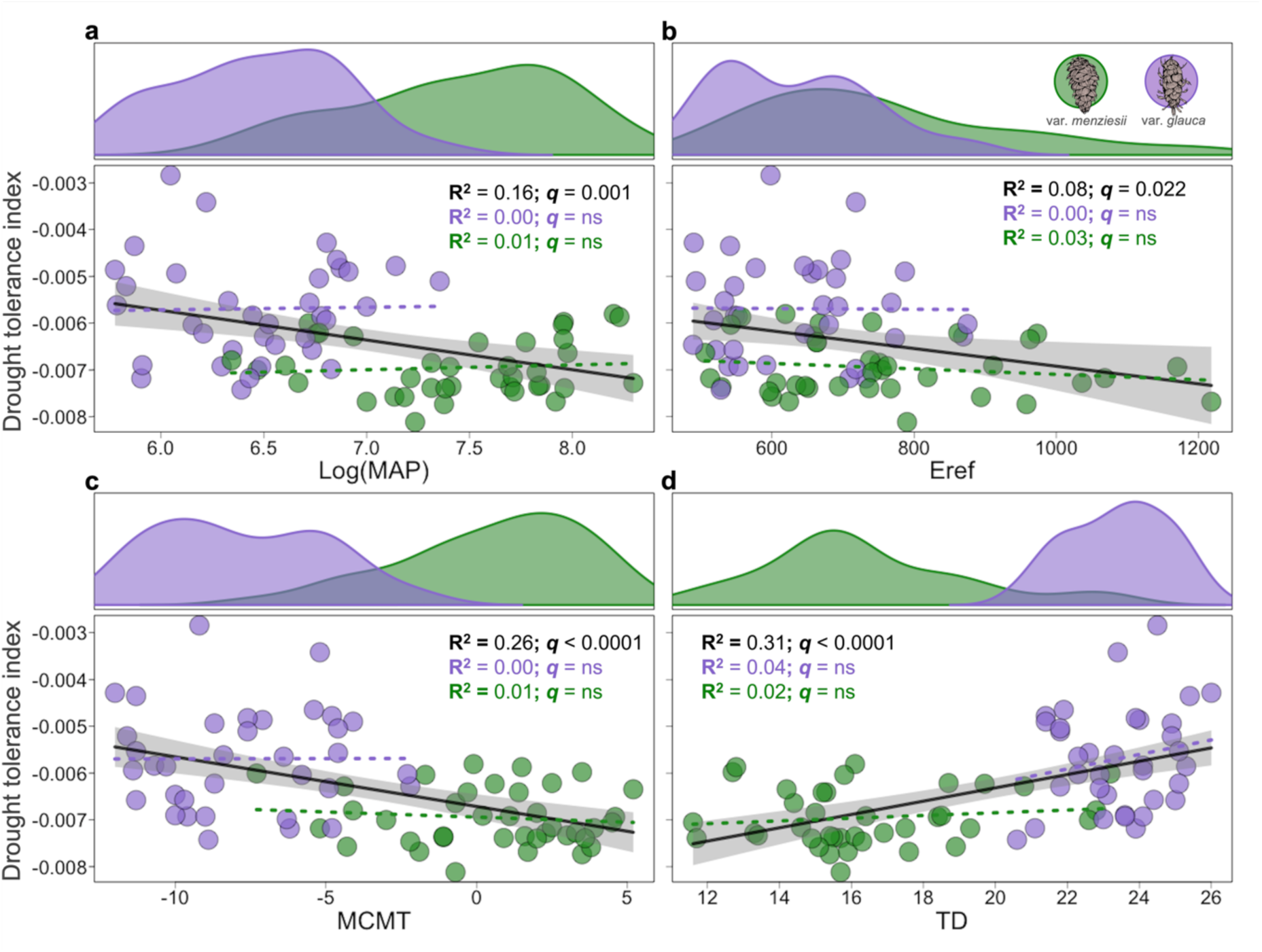
Relationships between drought tolerance index (DTI) and climatic gradients. (**a**) DTI regressed on mean annual precipitation (log (MAP (mm))). (**b**) DTI regressed on Hargreaves reference evaporation (Eref). (c) DTI regressed on mean coldest month temperature (MCMT (°C)). (**d**) DTI regressed on continentality (TD (°C)). The regression lines fitted at the species level (all sampled provenances) are depicted in black. 95% confidence intervals of significant relationships (*q-value* < 0.05) are depicted as ribbons. *q-value* is the same as the false-discovery-rate-adjusted *p-value* to account for multiple comparisons. Dashed lines show non-significant relationships (*q-value* > 0.05) within var. *glauca* (purple) and var. *menziesii* (green). Density plots depict the climate space of var. *menziesii* and var. *glauca*.

Despite the observed differentiation among provenances (V_POP-prov_) within var. *glauca* for DTI (Table 1), no significant phenotypic clines of drought tolerance along climate gradients were observed within this variety after correction for multiple comparisons. Within var. *menziesii*, consistent with the nearly absent signal of local adaptation to drought observed with the V_POP-prov_ analysis (Table 1), there were no significant relationships between geographic or climatic variables and DTI after correction for multiple comparisons (Table S5).

Significant clines were observed for THI species-wide and within var. *menziesii* under both drought and control treatments and across most tested climatic variables (Fig S10; Table S6), consistent with V_POP-prov_ estimates for drought and control treatments species-wide and within var. *menziesii* (Table 1). The THI_control_ associations with climate were stronger than the THI_drought_ associations across most of the climatic spaces occupied by var. *menziesii* (Table S6). Within var. *glauca*, despite the significant difference between THI_control_ and THI_drought_ (Fig. 2), no significant relationship was observed between THI and the tested climatic variables under both experimental conditions after correction of the *p-value* for multiple comparisons (Table S6).

### Plasticity and height growth-drought tolerance trade-off

Plasticity, as estimated by the relative height increment BLUEs of provenances in the control treatment minus the drought treatment was much greater on average in var. *menziesii* (0.18) than var. *glauca* (0.06) (Fig. 4c). There was a tendency for provenances within the range of var. *menziesii* that originated from more favorable environments (i.e., warmer and wetter) in western WA and OR to show higher degrees of plasticity relative to provenances from less favorable conditions (Fig. 4d; Fig S10), supporting the evidence of large variation for plasticity within this variety.

**Fig. 4.**
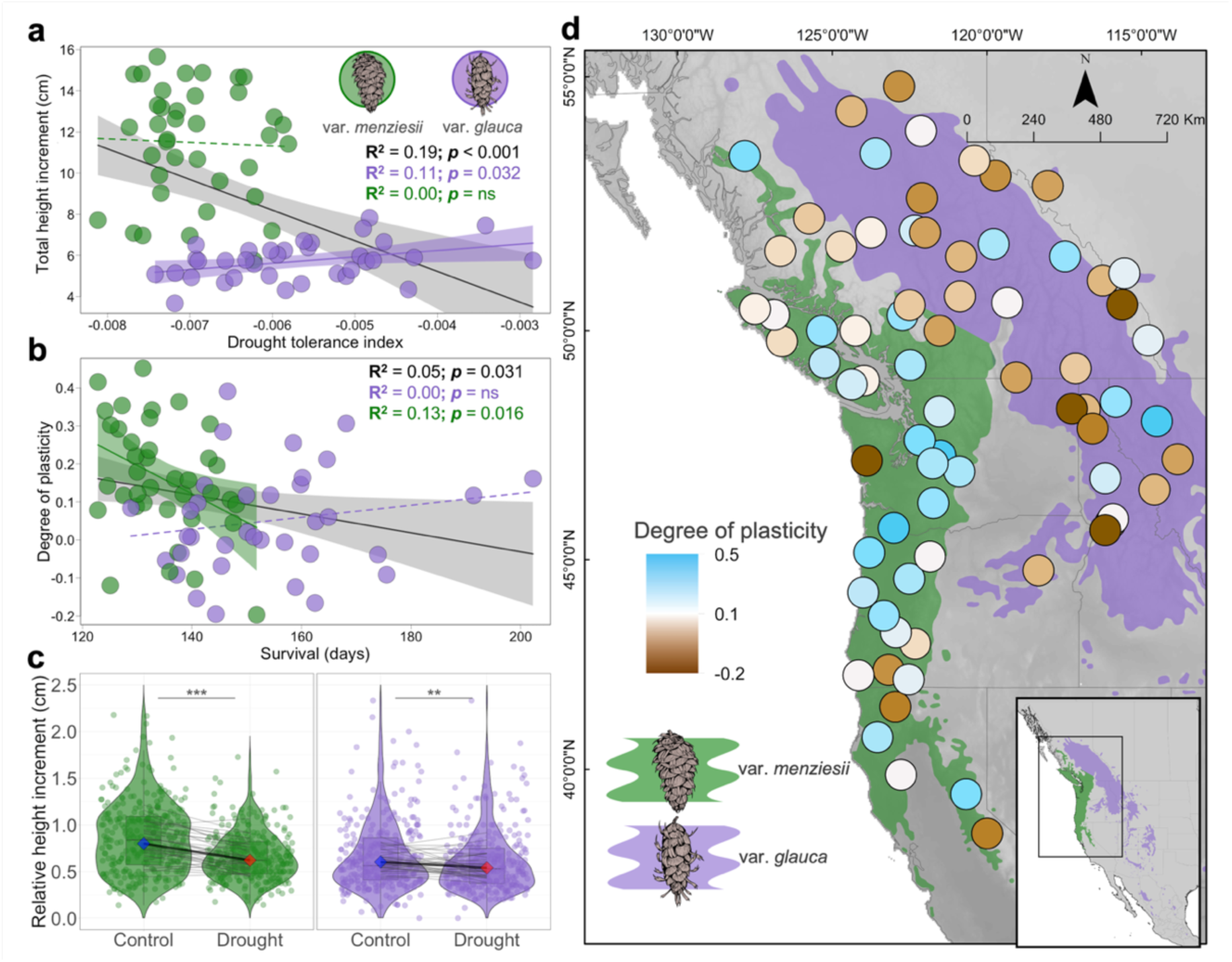
Height growth-drought tolerance trade-off and plastic response to drought in Douglas-fir. (**a**) Relationships between drought tolerance index (DTI) BLUEs in the drought treatment and total height increment (cm) BLUEs in the control species-wide and within varieties. (**b**) Relationships between survival BLUEs (estimated as the number of days in the experiment until seedlings reach the F_v_/F_m_ threshold of 0.4) and plasticity (the estimated relative height increment BLUEs in the control treatment minus the estimated BLUEs in the drought treatment) species-wide and within varieties. (**c**) Relative height increment in drought and control treatments and the reaction norms (black lines) for each variety. Blue and red diamonds, BLUEs estimated for control and drought treatments, respectively, within each variety. The plasticity between treatments within each variety was tested with the non-parametric Wilcoxon Test. Shaded grey lines depict provenance reaction norms. (d) Species-wide distribution of plasticity. Significant regression lines (*P* < 0.05) in **a** and **b**, are depicted in black for the species-wide analysis, in purple for var. *glauca*, and in green for var. *menziesii*. Non-significant relationships are depicted as dashed lines. Confidence intervals of significant relationships are depicted as ribbons. **, *p* < 0.01; ***, *p* < 0.001; ns, *p* > 0.05.

Species wide, a negative and significant relationship between growth and drought tolerance was detected (for THI versus DTI, R^2^ = 0.19, *p* < 0.0001 in the control treatment, and R^2^ = 0.12, *p* = 0.0026 in the drought treatment), indicating a trade-off between growth and drought tolerance (Fig. 4a; Fig. S3). However, this relationship was largely driven by differences between varieties. No significant relationship was detected between growth and drought tolerance within var. *menziesii*, and a weak but significant positive relationship was observed within var. *glauca* (R^2^ = 0.11, *p* = 0.032 in the control treatment, and R^2^ = 0.29, *p* = 0.0008 in the drought treatment).

## Discussion

### Strong differences between varieties dominate provenance variation in drought tolerance across the species

Douglas-fir is known for its ability to tolerate dry conditions (Huang et al., 2022; Stangler et al., 2022) and being relatively resilient to unfavorable drought periods that restrict growth (Song et al., 2022; Vitali et al., 2017). Consistent with this, we observed only a slight reduction in photosynthetic efficiency one month after the start of the drought treatment in the experiment. This treatment effect increased substantially in every subsequent measurement, but it was only after 71 days under extreme drought that seedlings showed more than 50% reduction in F_v_/F_m_ on average. At this level of reduction, photosynthetic tissues are likely permanently damaged (Brodribb et al., 2021) and, in general, plants can be considered dead, with exceptions (e.g., Woo et al., 2008).

In the species-wide analyses of all provenances tested, we observed a weak signal of local adaptation to drought in seedlings, evidenced by provenance variation and clines along temperature and, to a lesser extent, precipitation-related gradients in F_v_/F_m_ measurements on individual dates as well as in the drought tolerance index (DTI, the rate of decline in F_v_/F_m_) (Fig S5; Fig. 3). The extent of provenance differentiation in F_v_/F_m_ varied as drought progressed (Table 1). It primarily resulted from the large difference between var. *glauca* and var. *menziesii* provenances in the timing of responses to the drought (Table 1). Considerable differences in the climate niches and selection pressures experienced by each variety since their divergence, circa 2.1 Mya (Gugger et al., 2010), help explain the large phenotypic differences observed between them and the resulting species-wide relationships between drought tolerance and provenance climate (Fig. 3). For example, the slower response to drought in var. *glauca* seedlings reflects adaptation to the generally harsher continental climate on the eastern and northern portions of the species range, with drier summers as well as colder and longer winters. Variety *menziesii*, on the other hand, is adapted to mild, wetter maritime climates and long frost-free periods within most of the coastal region of the Pacific Northwest – conditions favorable for highly productive and competitive forests (Howe et al., 2006). Adaptations such as cold acclimation and earlier bud set to escape early frost in the fall are expected to increase survival and fitness under more stressful climatic conditions while restricting growth (Rehfeldt, 1977). These adaptations may be correlated with drought tolerance, given that some morpho-physiological mechanisms used by conifers to tolerate freezing and drought stresses are expected to be the same or related (Bansal et al., 2016; White, 1987). However, in a parallel genome-wide association study of a subset of the provenances included here, we found little overlap between genes associated with cold hardiness and drought tolerance (R. Candido-Ribeiro et al., University of British Columbia, *unpublished data*). We detected stronger signals of local adaptation for height growth than for drought tolerance (Table 1; Table S5; Table S6). We observed a shorter window of primary growth in var. *glauca* than var. *menziesii*, regardless of treatment (drought or control) or region of origin, with most var. *glauca* individuals ceasing or slowing height growth well ahead of var. *menziesii* seedlings and sustaining less mortality under severe drought (Fig. 4b; Fig. S12). The faster growth of var. *menziesii* has been observed elsewhere (Rehfeldt et al., 2014). Provenances of many other temperate conifer species show a general pattern of greater growth, lower cold hardiness, and delayed growth cessation in provenances from warmer environments (Alberto et al., 2013; Aitken and Bemmels, 2016; Rehfeldt et al., 2018). These studies have consistently found that for temperate trees, cold temperatures limit growth, and indicate generally that local adaptation to climate is driven more strongly by temperature than by precipitation related variables (Leites & Benito Garzón, 2023). Temperature related variables were also found to have, in general, the strongest associations with radial growth of mature trees in common gardens in response to drought (Montwé et al., 2016; Sang et al., 2019). This may be because temperature in temperate and boreal regions varies seasonally at broader and more predictable time and geographic scales than precipitation (Samset et al., 2019), providing more consistent conditions for divergent selection.

Plastic responses may also play an important role for populations under increasingly variable climates and more extreme weather events (Alberto et al., 2013). Here we observed that var. *glauca* provenances show little plasticity for growth between well-watered and drought conditions. In contrast, var. *menziesii* provenances were much more plastic, with trees originating from the most productive forest regions showing the largest differences in growth between drought and control treatments (Fig. 4d). This growth plasticity in var. *menziesii* likely evolved as a response to inter- and intra-specific competition for light and other resources in highly productive coastal rainforests. However, these plastic coastal provenances appear to lack sufficient adaptive variation to withstand extreme drought events, perhaps lacking detection of cues to cease growth and prepare physiologically and developmentally for stress as observed in var. *glauca*. Therefore, under increasing severe drought, this plasticity may become maladaptive (Fig 4b).

### Most variation in drought tolerance within varieties is within provenances

Provenance differentiation within varieties for individual F_v_/F_m_ measurements was surprisingly weak. Within var. *menziesii*, we only observed V_POP_ estimates higher than F_ST_ (Table 1) during the first three of five individual F_v_/F_m_ measurements, when drought was either not present or was moderate. In contrast, within var. *glauca*, provenance differentiation for F_v_/F_m_ was highest for the last three measurements (moderate to extreme drought), when V_POP_ surpassed F_ST_ (Table 1). While some var. *glauca* provenances could maintain their photosynthetic capacity under extreme drought (e.g., provenances 4, 38, and 48 within the northernmost region), others had largely succumbed (e.g., provenances 3 and 34 within southern British Columbia). This variation produced weak population differentiation for the drought tolerance index (DTI) within var. *glauca* (V_POP_ = 0.13). The strongest adaptive differentiation among provenances somewhat surprisingly was in the northernmost provenances in British Columbia, the coldest but not the driest region.

Within var. *menziesii*, differentiation for DTI was absent (V_POP_ = 0.01). This result contrasts with the significant differences among provenances found by Bansal et al. (2015a) for traits presumably related to drought tolerance (i.e., minimum transpiration, water deficit and specific leaf area), and relationships between these traits and climate of origin of a similar number of coastal Douglas-fir provenances sampled in our study. Rather than impose an experimental drought treatment, they established common garden experiments at three test sites varying in temperature and moisture, and seedlings at the hottest, driest site did not experience conditions as stressful as in our experiment. All traits in Bansal et al. (2015a) were associated with growth (height or diameter), while minimum transpiration was also associated with cold hardiness (Bansal et al., 2015b), and the most relevant environmental drivers of adaptive differentiation in water relations traits were minimum temperature and elevation. Differences in the study designs could partially explain the contrasts between their results and ours, as we targeted provenances relatively distant from each other and sampled across a larger area. Additionally, we assessed traits that indicated a direct physiological response to the drought stress. Interestingly, we observed a lack of trade-off between drought tolerance and height growth within varieties (Fig. 4a), which also, to some extent, contrasts with Bansal et al.’ (2015a) results.

The weak to non-existent local adaptation to drought in seedlings within varieties and the maintenance of standing variation within provenances has several potential and non-exclusive explanations. First, gene flow levels are high for this and other wind-pollinated trees (Kremer et al., 2012). If selection, i.e., drought, varies weakly at large spatial scales and is not sufficiently strong or consistent over time to overcome the homogenizing effects of gene flow, local adaptation is expected to be weak or absent (Yeaman, 2022). Local adaptation is less constrained by gene flow when traits are controlled by few loci of large effects (Yeaman, 2022); however, drought tolerance in Douglas-fir is highly polygenic (R. Candido-Ribeiro et al., University of British Columbia, *unpublished data*). Furthermore, temporal variation in drought could maintain high within-population genetic variation (Cavender-Bares, 2019).

Second, if adaptive differentiation has occurred at micro- or meso-geographic scales, as has been observed for some other conifers (Budde et al., 2023; Scotti et al., 2023) including coastal Douglas-fir (Campbell, 1979), the widescale sampling in this project may not have detected this variation. Water availability and soil drought stress can vary on a fine scale, where local topographic and edaphic features produce fine grained spatial variation of selection (Ferrell & Woodard, 1966). Thus, drought stress operating as a small-scale selective force, countered by the effects of gene flow, could have contributed to the high within-provenance variation and the lack of differentiation among sampled provenances within varieties. Future drought studies combining range-wide and localized sampling designs could produce a better understanding on the scale at which different traits have locally adapted in conifer species.

Third, responses to drought in conifers are complex and likely involve different traits at different stress levels, seasonal timing and duration, and these traits may vary in importance among regions (Moran et al., 2017). This makes trait and phenotyping method selection challenging. Among commonly used physiological traits for assessing responses of tree seedlings to drought, F_v_/F_m_ is one of the last to exhibit substantial changes to a point of no return, but is also in general more variable than, for example, the thresholds for the percentage loss of rehydration capacity (Trueba et al., 2019). We found individual F_v_/F_m_ measurements were useful for characterizing drought tolerance within varieties and provenances but required assessments on multiple dates over time to be informative. Our drought tolerance index (DTI), describing the linear decline in photosynthetic efficiency of plants under drought stress, was useful for assessing and comparing large numbers of seedlings and has been shown to predict seedling mortality, but might be limited in identifying more nuanced differences (i.e., within varieties). Advancement of image processing for high-throughput phenotyping in plants (e.g., Exposito-Alonso et al., 2018) could improve and optimize assessments of drought tolerance in trees substantially.

Finally, seedlings represent an important and vulnerable life history stage of trees experiencing increasing climatic stresses. Seedlings cohorts typically possess large amounts of adaptive genetic variation on which selection can act. However, traits responding to selection in seedlings might not be the same as traits responding to the same selective pressures in older trees, and local adaptation in adult cohorts is likely to reflect multiple waves of selection that populations have endured over the course of their life spans. Thus, the patterns of local adaptation to drought observed in seedlings may differ from those observed in adults. In fact, multiple range-wide genecology studies with widespread conifers have identified strong signals of local adaptation in traits related to drought in older trees (Montwé et al., 2015; Montwé et al., 2016; Sang et al., 2019). Therefore, the potential ontogenetic changes in the patterns and relationships of drought tolerance represent another limitation of our study. Future studies should evaluate whether more drought tolerant seedlings maintain their fitness advantage in responding to drought at later stages, but such long-term studies are rare because they are difficult.

### Mitigating the effects of climate change

With extreme weather events becoming increasingly frequent and severe in the Pacific Northwest due to anthropogenic climate change (e.g., White et al., 2023), drought is expected to play a progressively larger role in driving selection in these ecosystems (van Mantgem et al., 2009; Yuan et al., 2023). Mismatches between historically locally adapted phenological patterns and rapidly changing climates are, therefore, expected to result in population maladaptation (Aitken et al., 2008; St Clair & Howe, 2007). Assisted gene flow (AGF) has been proposed for temperate tree species as a potential strategy to maintain growth potential (i.e., fitness) of local populations under future warmer climates (Aitken & Bemmels, 2016; Aitken & Whitlock, 2013). However, the weak signal of local adaptation to drought observed in our study suggests that AGF will be less effective for mitigating the effects of increasing summer droughts on seedlings than for facilitating adaptation to rising temperatures within Douglas-fir varieties.

On the other hand, the high level of phenotypic variation in drought tolerance within provenances relative to that observed among provenances suggests considerable standing genetic variation that could be used, through natural selection or breeding, to increase drought tolerance under climate change. Given the long time required for natural populations of conifers to adapt to environmental changes (Aitken et al., 2008; Aitken & Bemmels, 2016), selective breeding for drought tolerance may be an important tool for tackling the ongoing effects of climate change within varieties. The narrow-sense heritability of DTI, estimated as approximately 0.2 in a var. *menziesii* breeding population in a follow-up experiment (Nuhu, 2022), shows promise for selecting drought tolerant families within provenances. Furthermore, the lack of trade-offs between drought tolerance and growth suggests that selecting for greater drought tolerance within varieties will not compromise height growth in seedlings. Surprisingly, some more tolerant provenances within var. *glauca* had greater height growth under the severe drought conditions imposed in this study than under the control treatment. Thus, it may be possible to increase both growth and drought tolerance through selection.

Lastly, besides selective breeding, silvicultural decisions are likely to play a progressively larger role in boosting seedling survival and establishment under increasing drought conditions in the Pacific Northwest. Practices such as careful selection of species (MacKenzie & Mahony, 2021), larger root plugs to improve root development, and seedling drought hardening prior to planting (Grossnickle, 2012), as well as site and microsite selection and preparation, will be crucial for planted seedlings from selected families to express their phenotypes and provide forest ecosystem services.

## Acknowledgement

This work was part of the CoAdapTree project led by S.N.A and majorly founded by Genome Canada (241REF; Co-Project Leaders Sam Yeaman and Richard Hamelin), with co-funding from Genome BC and 16 other sponsors (http://coadaptree.forestry.ubc.ca/sponsors/), including Genome Alberta, Génome Québec, the British Columbia Ministry of Forests, Lands and Natural Resource Operations (BCMFLNRO), Canadian Forest Service (Natural Resources), Alberta Innovates Bio Solutions, Vernon Seed Orchard Company, University of Alberta, University of British Columbia, the Forest Genetics Council of British Columbia, Compute Canada, Mosaic Forest Management, TimberWest, and Western Forest Products. Douglas-fir seeds were kindly donated by 16 forest companies and agencies in Canada, United States, and Mexico (visit https://coadaptree.forestry.ubc.ca/seed-contributors/ for the names of all contributors). We thank for the invaluable technical assistance and support from many current and former members of the Aitken lab at the University of British Columbia, without whom this study would not have been made possible. Particularly Christine Chourmouzis and Pia Smets for helping with the selection and acquisition of seedlots, seed stratification and sowing, and assisting at all stages of the experimental establishment and data collection. We thank Dragana O. Vidakovic and Joanne Tuytel for their assistance in establishing the experiment. Alex Girard helped extensively and tirelessly with data collection and maintenance of the experiment. Martin Henry kindly provided the Douglas-fir cone drawings used in the figures. Rob Guy provided helpful suggestions on the experimental design. Jon Degner gave helpful insights on the analysis. Thanks to Hayley Tumas and Beth Roskilly for their comments and suggestions on this manuscript.

## Author contributions

SNA and RCR designed the study. RCR conducted the experiment, collected the data, and performed the analyses with input from SNA. Both authors contributed to the writing of the manuscript.

## SUPPORTING INFORMATION

### Supplementary Text

#### Pre-experimental steps

Seeds were stored at 4°C until stratification (24 h submerged in aerated water followed by 28 days at 4°C to break embryo dormancy). In May 2017, two seeds were sowed per individual container (55 cm^3^ *Stuewe & Sons* Ray Leach UV, with commercial substrate). If two seedlings germinated, one was removed. The seedlings were kept in trays of 200 plants each under controlled environmental conditions in the UBC Botany Greenhouse at the University of British Columbia, Vancouver. They were well-watered, fertilized with an N-P-K solution 11-41-8 + micronutrients, and kept at temperatures between 20°C and 25°C for approximately five months. The trays were rotated every three days to avoid greenhouse position effects until the plants were transplanted into the experiment. In order to induce bud set and overwinter the plants, the seedlings were then exposed to outdoor conditions (chilling temperatures) and limited daylight (∼ 9 – 10 h) for six weeks.

#### Correction for spatial autocorrelation of drought effects and estimation of DTI (drought tolerance index)

An autoregressive model of residuals was fit for each F_v_/F_m_ measurement date in the package *ASReml-R* 4.0 (Butler, et al., 2017) to reduce spatial pattern effects of drought within boxes. F_v_/F_m_ was the dependent variable, and a single fixed intercept was estimated in each model. Row and column positions within experimental blocks were modeled as random residual effects with a correlation structure. The residuals were assumed to be the spatial effect on each plant and were, therefore, subtracted from the F_v_/F_m_ values (Fig. S1). This step was only applied to the drought treatment as no spatial effect in drought response is expected in the control treatment.

The corrected F_v_/F_m_ values from each individual were then regressed on the days of the experiment. The following linear model was used to estimate the rate of decline in F_v_/F_m_ per day under drought conditions (DTI):

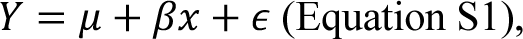

where *Y* is the F_v_/F_m_ (after correction for spatial autocorrelation), 𝜇 is the estimated intercept, 𝛽 is the estimated effect (slope) of days in the experiment (*x*) on *Y* and 𝜖 is the error term. The slope of each linear regression is the rate of decline in chlorophyll fluorescence of each plant per day (DTI) and was used as a proxy for drought tolerance of each individual.

### Estimating provenance BLUEs before testing for associations with the environment

For the estimation of BLUEs, the following mixed-effects model implemented in *ASReml-R* 4.0 (Butler, et al., 2017) was used:

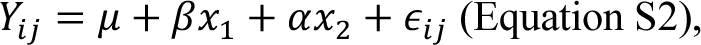

where 𝑌_*ij*_ is the phenotype corresponding to individuals from provenance *i* and block *j*; 𝜇 is the phenotype global mean across all individuals within one of the treatments (fixed intercept), 𝛽 is the coefficient for the fixed effect of provenance (𝑥_$_) and 𝛼 is the random effect of blocks (𝑥_%_) and 𝜖_*ij*_ is the error term for 𝑌_*ij*_. For total height increment of drought and control treatments, rows and columns were incorporated into the residual term of the model with a correlation structure to account for possible effects of position within the experiment on the phenotype. We did not account for the effect of position on the estimation of BLUEs for DTIs to avoid over correction, given that individual F_v_/F_m_ measurements were already corrected. When necessary, we first used an arcsine square root transformation of DTIs (after adding a constant to make them positive) and F_v_/F_m_ measurements, and a square root log transformation of total height increment and relative height increment to meet the assumptions of normality of residuals and homoscedasticity of variances. BLUEs were estimated with the function *predict* in *ASreml-R* (Butler et al., 2017).

## Supplementary Figures

**Fig. S1.**
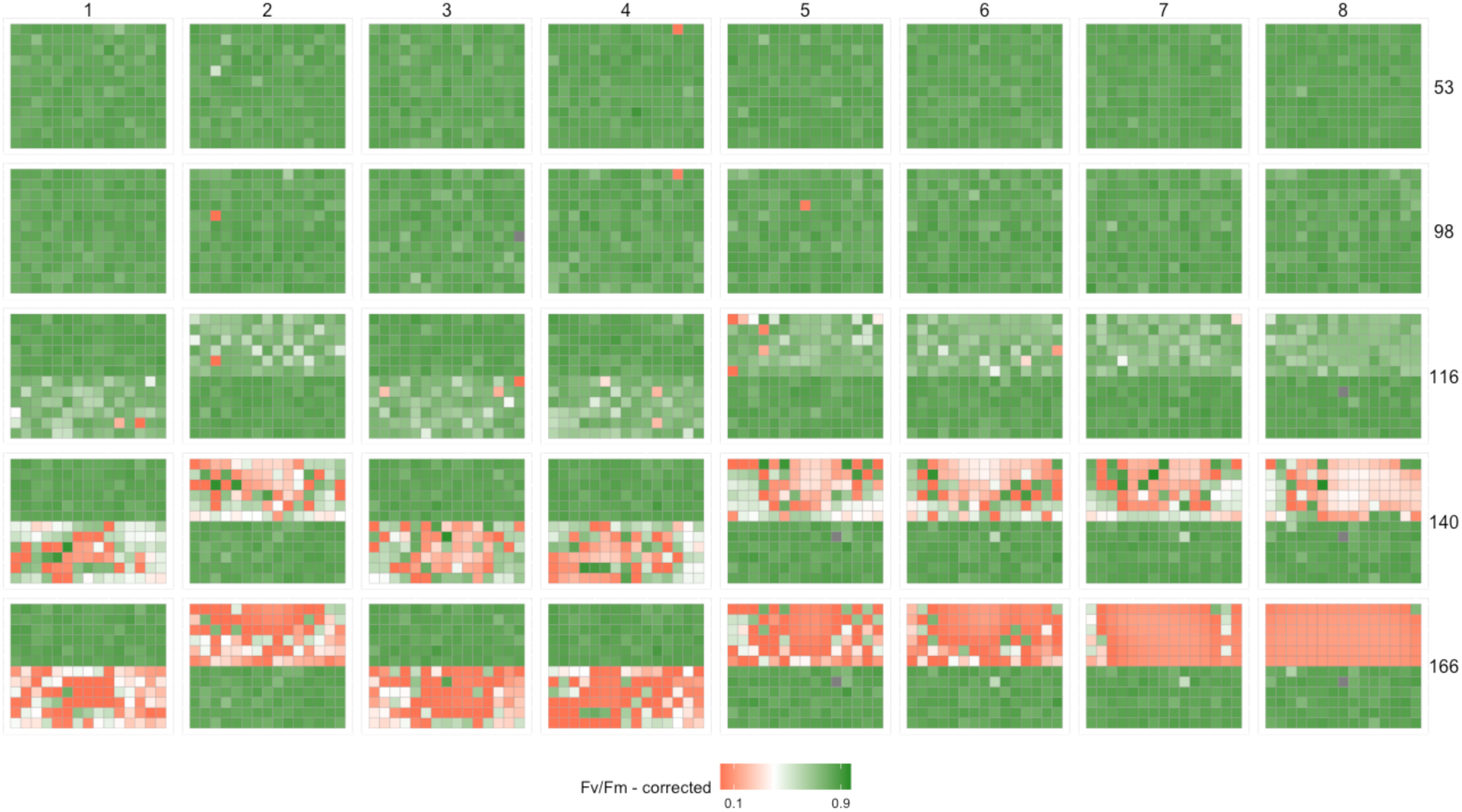
Chlorophyll fluorescence (F_v_/F_m_) data corrected for spatial autocorrelation in the experiment through time. Major columns depict blocks 1 through 8, and major rows depict the day of experiment (53 to 166 days) when F_v_/F_m_ was measured. Each block is represented by one box (subplot) with a drought to death treatment (with a progressing effect on F_v_/F_m_) and an adjacent control box (mostly stable F_v_/F_m_ values through time). Each square within a block is an individual plant. Edge trees around each box are not shown. Only F_v_/F_m_ obtained from plants under the drought treatment were corrected for spatial autocorrelation.

**Fig. S2.**
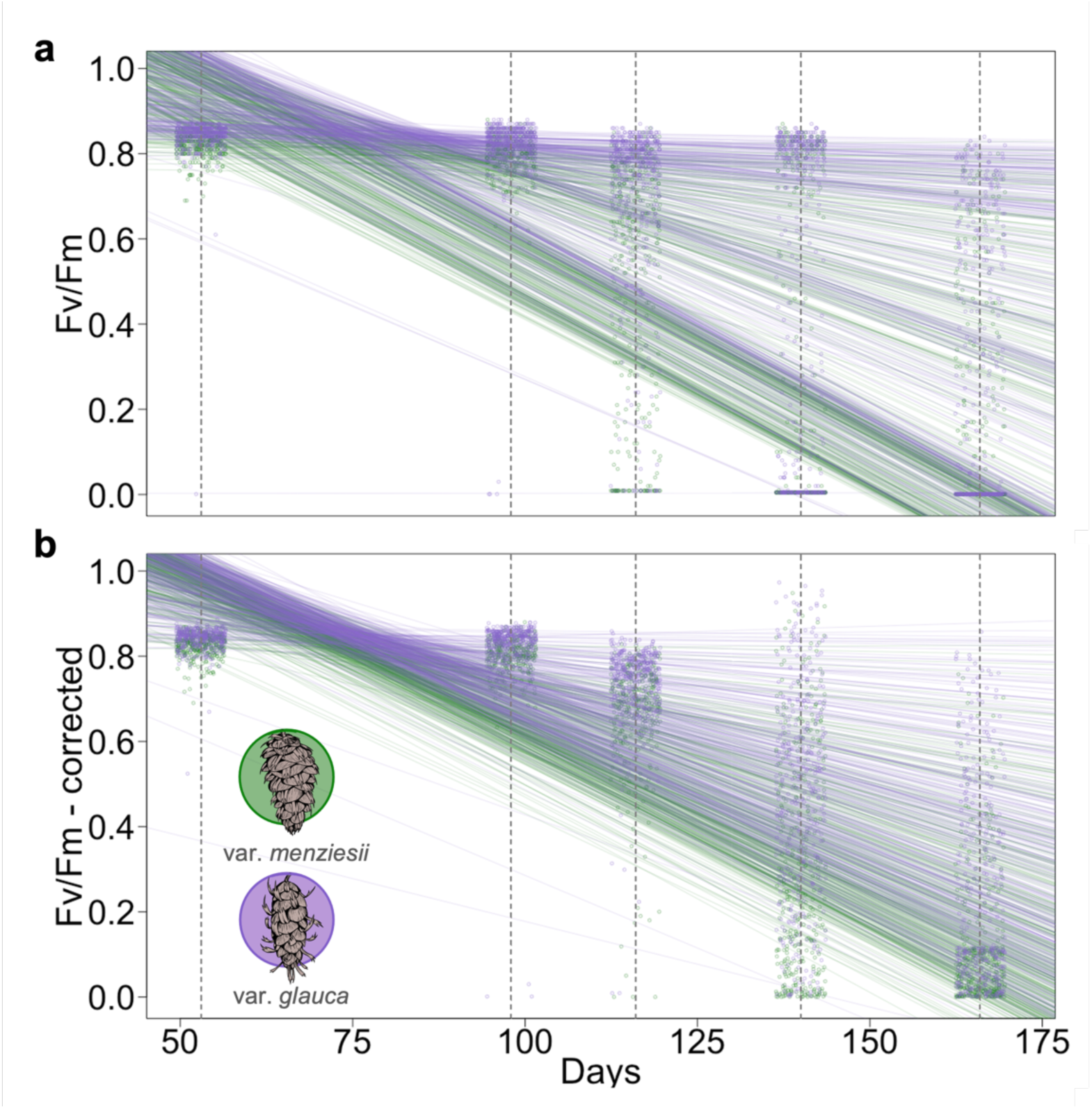
Individual seedling progress of F_v_/F_m_ (chlorophyll fluorescence) in the drought treatment. (**a**) F_v_/F_m_ uncorrected and (**b**) corrected for spatial autocorrelation. F_v_/F_m_ values were corrected with an autoregressive model of residuals implemented in the package *ASReml-R* 4.0 (Butler et al., 2017). Lines were fitted with a simple linear regression of F_v_/F_m_ values regressed on the day of experiment. The slope of each simple linear regression in **b** represents the individual DTI (drought tolerance index).

**Fig. S3.**
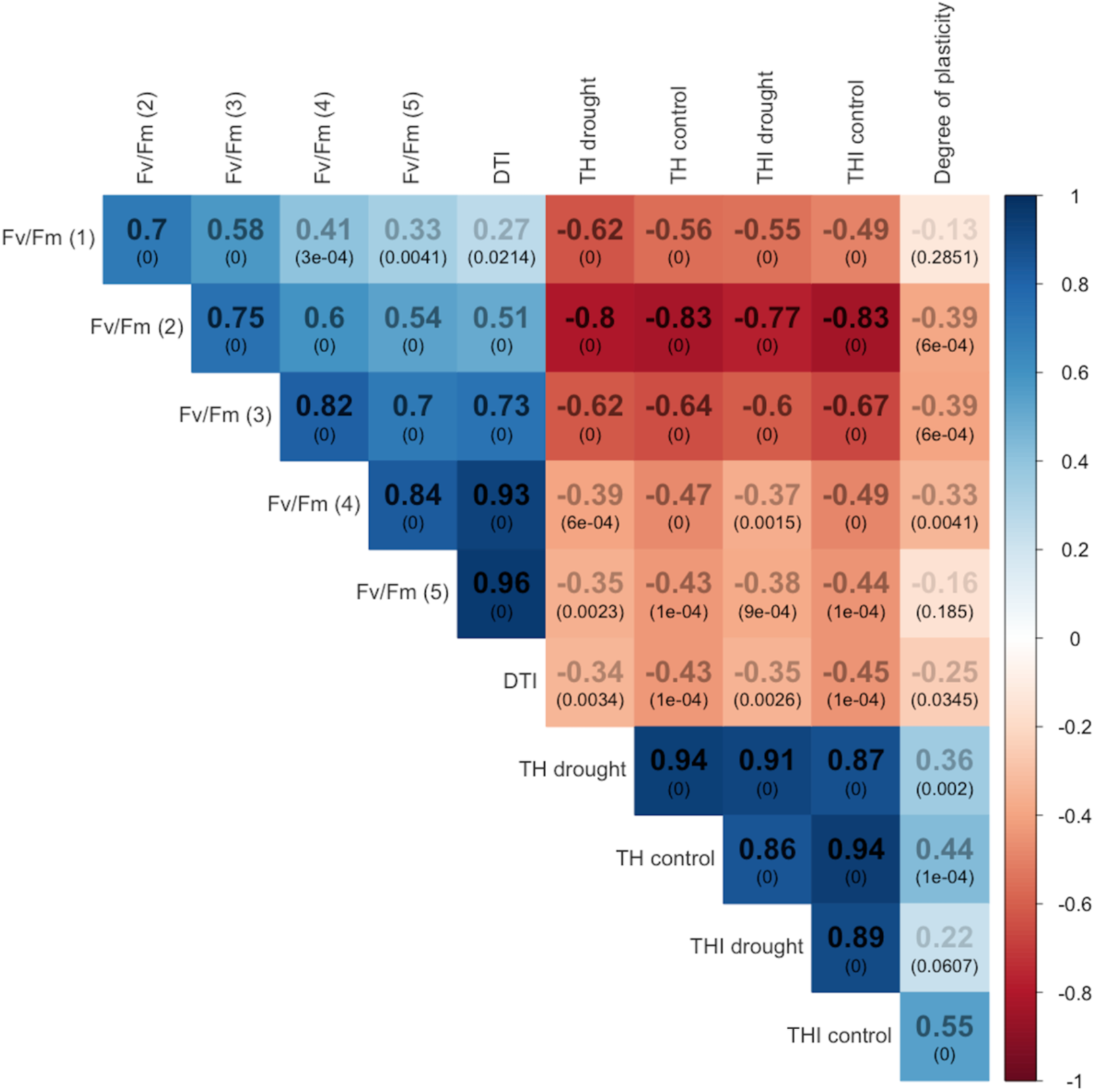
Correlations between all estimated trait BLUEs for provenances species wide (with the two varieties included). The heatmap depicts the Pearson’s correlation coefficient (*r*) between the trait BLUEs (in shades of black). F_v_/F_m_ (i), photosynthetic efficiency on individual date measurements. F_v_/F_m_ measurements and DTI are the provenance BLUEs estimated from the drought treatment. Measurement 1 was performed 16 days before the drought treatment started (53 days after the establishment of the experiment). Measurements 2, 3, 4, and 5 were performed 29, 47, 71, and 97 days respectively after the drought treatment started. *P-values* are shown in parentheses.

**Fig. S4.**
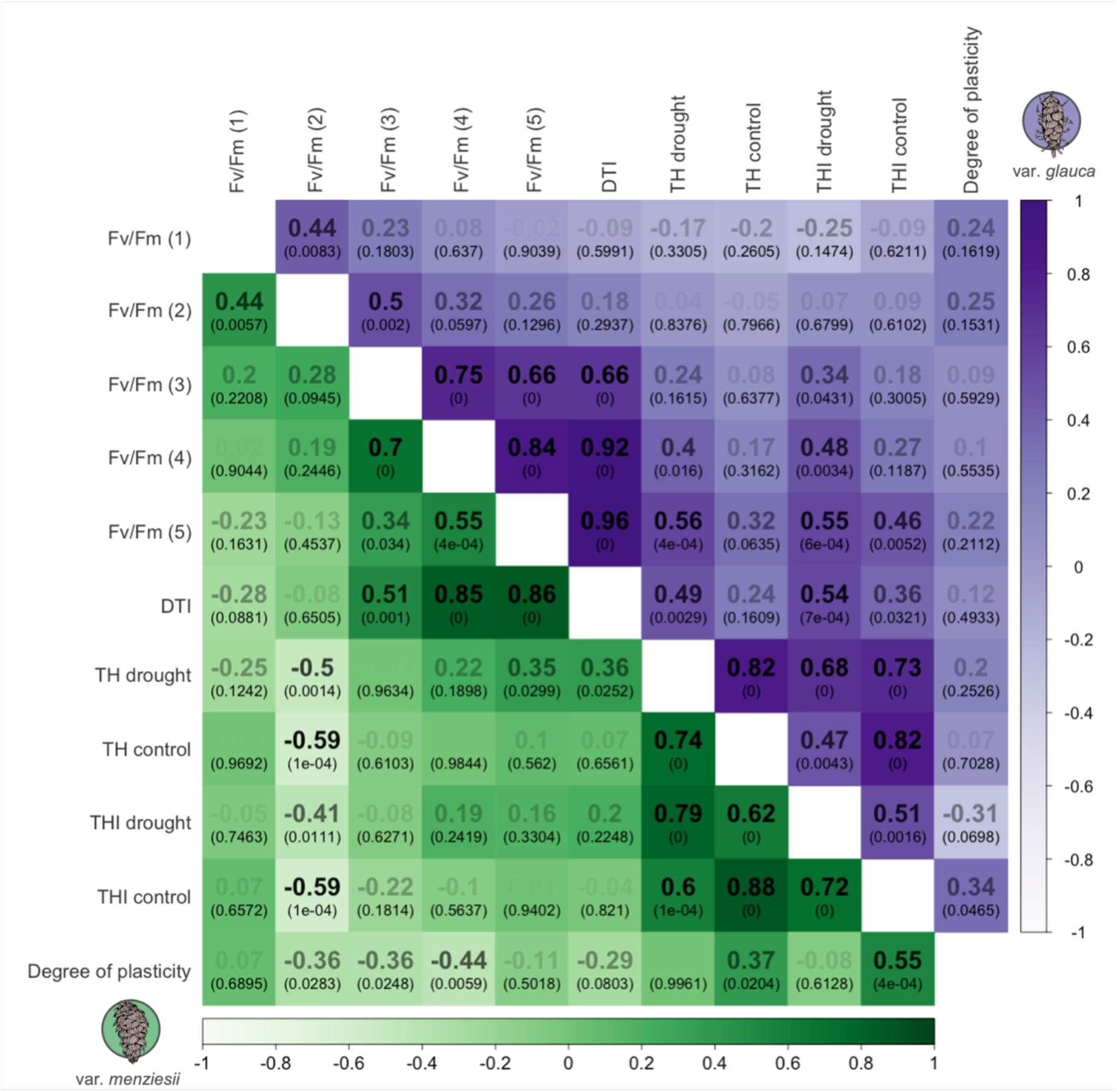
Correlations between all estimated trait BLUEs for provenances within varieties. The heatmap depicts the Pearson’s correlation coefficient (*r*) between the trait BLUEs (in shades of black). Associations depicted below the diagonal are var. *menziesii*’s, and above the diagonal var. *glauca*’s. F_v_/F_m_ (i), photosynthetic efficiency on individual date measurements. F_v_/F_m_ measurements and DTI are the provenance BLUEs estimated from the drought treatment. Measurement 1 was performed 16 days before the drought treatment started (53 days after the establishment of the experiment). Measurements 2, 3, 4, and 5 were performed 29, 47, 71, and 97 days respectively after the drought treatment started. *P-values* are shown in parentheses.

**Fig. S5.**
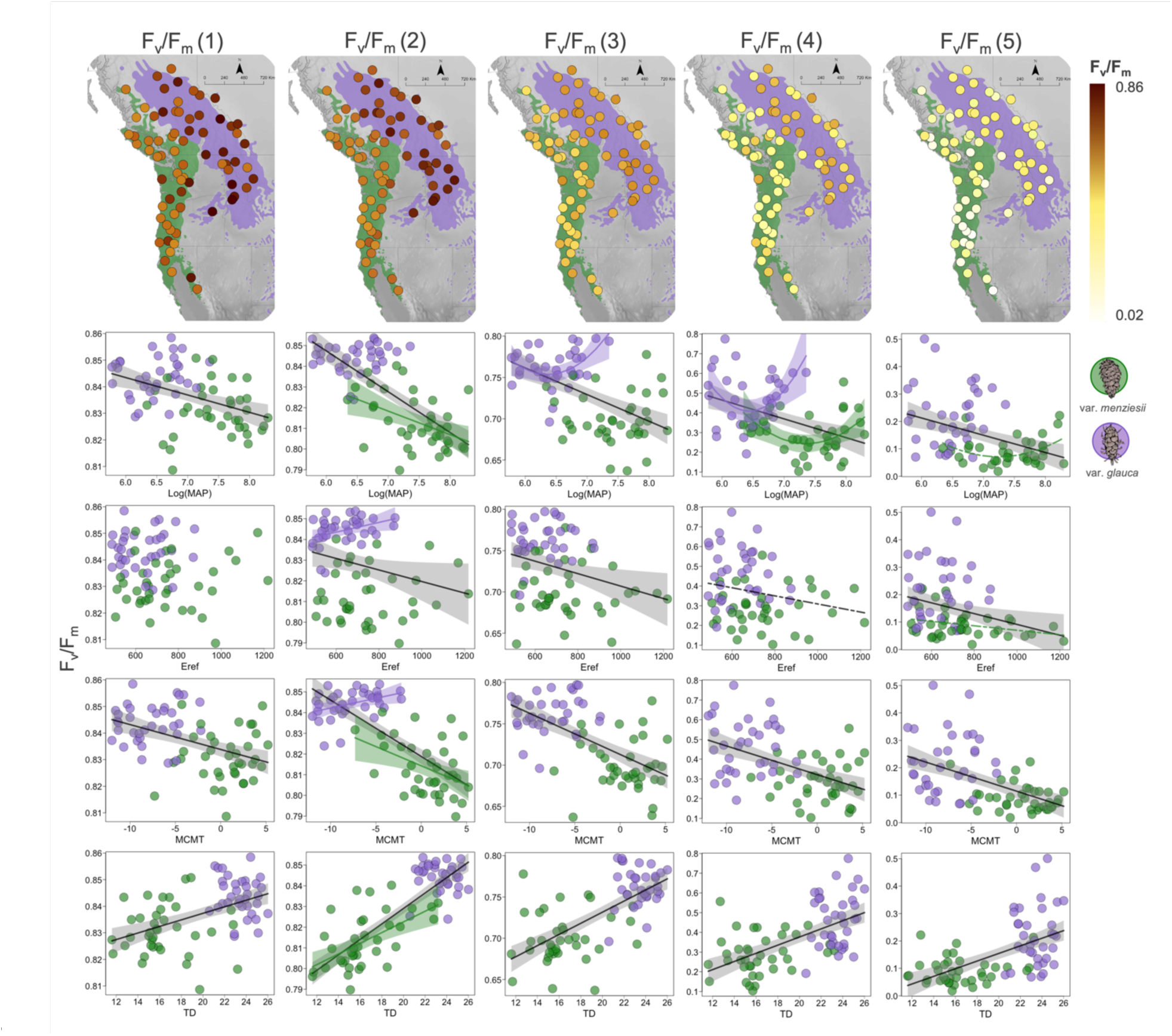
Provenance variation across the range and clines along climatic gradients for F_v_/F_m_ measured on individual dates in the drought treatment. F_v_/F_m_ (i), photosynthetic efficiency on individual date measurements. F_v_/F_m_ measurements were corrected for spatial autocorrelation. Measurement 1 was performed 16 days before the drought treatment started (53 days after the establishment of the experiment). Measurements 2, 3, 4, and 5 were performed 29, 47, 71, and 97 days respectively after the drought treatment started. Points in the maps and plots depict provenance BLUEs for F_v_/F_m_. See Table S7 for R^2^ and *p-values* of the F_v_/F_m_ vs. climate relationships.

**Fig. S6.**
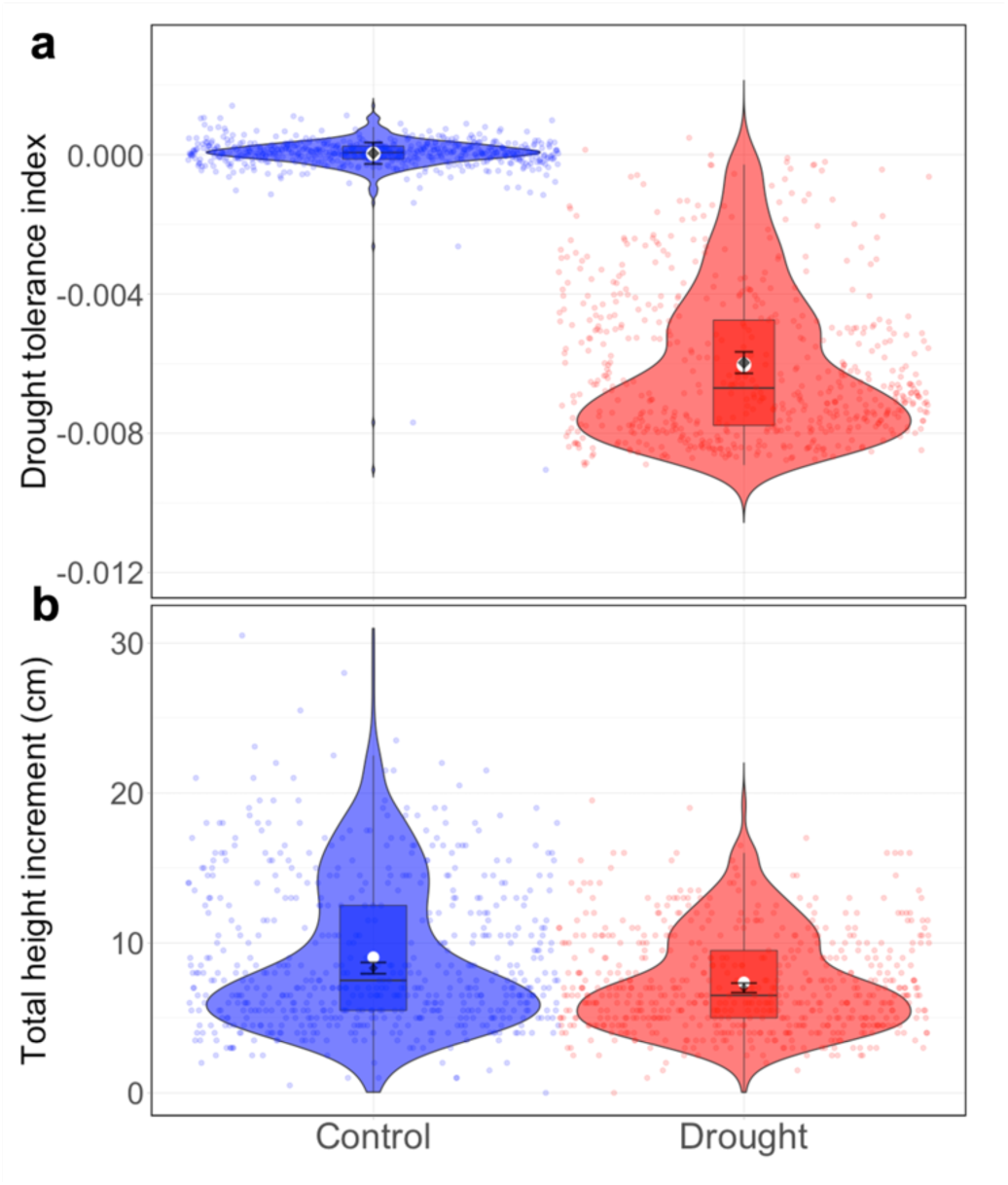
Species-wide effect of treatment (control vs. drought) on traits. Drought tolerance index (DTI) (**a**), and total height increment (THI) (**b**) for control (blue) and drought (red) treatments at the species-wide level (var. *menziesii* and var. *glauca* together). White dots depict the global averages for each treatment. Black diamonds and error bars show BLUE values and standard errors, respectively. See Table S2 for statistical test results.

**Fig. S7.**
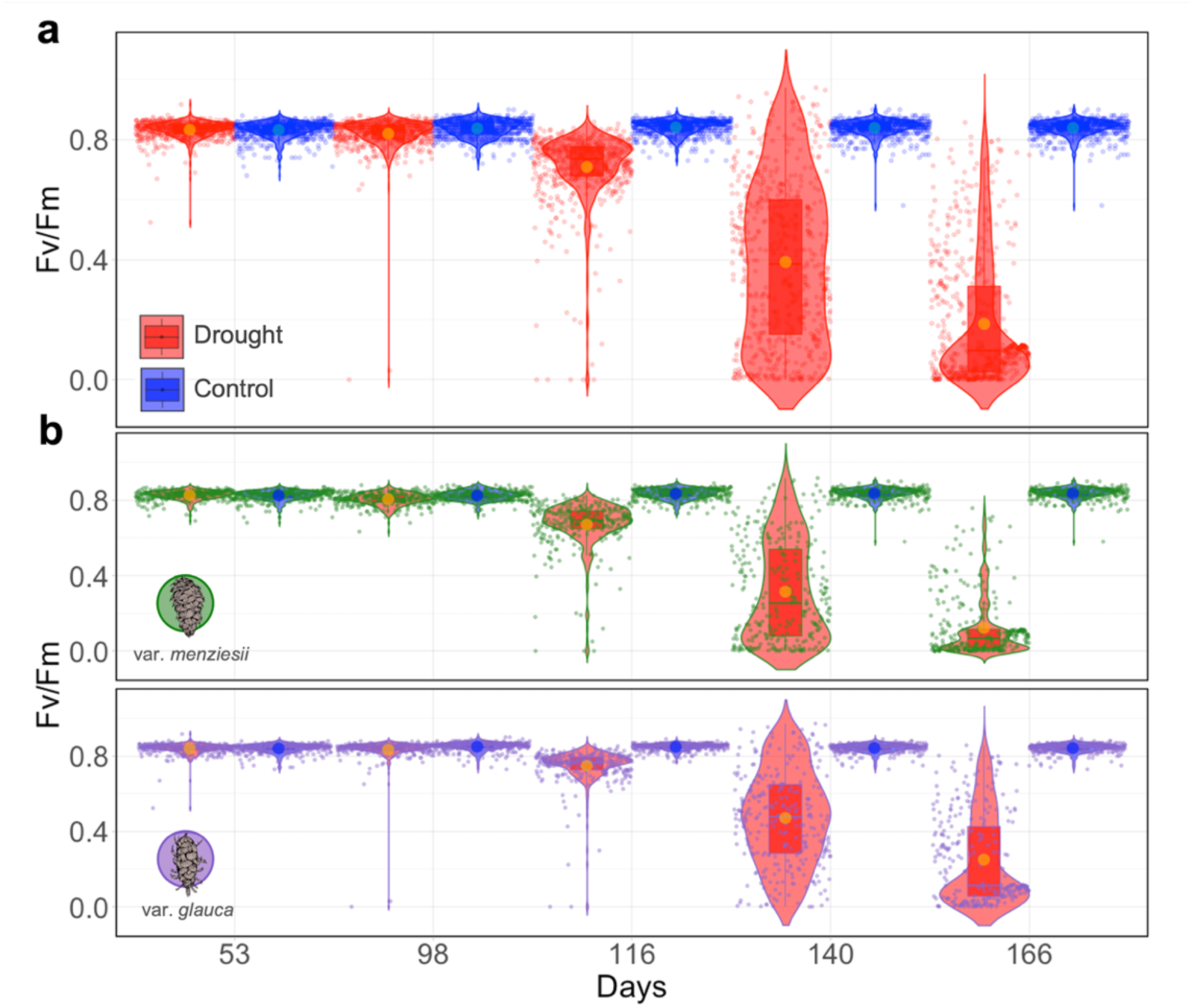
Effect of treatment (control vs. drought) on individual F_v_/F_m_ measurements. Species-wide (**a**) and within varieties (**b**) level testing. Individual F_v_/F_m_ measurements in the drought treatment were corrected for spatial autocorrelation. Orange and blue dots depict the F_v_/F_m_ averages for the drought and control treatments respectively. See Table S2 for statistical test results.

**Fig. S8.**
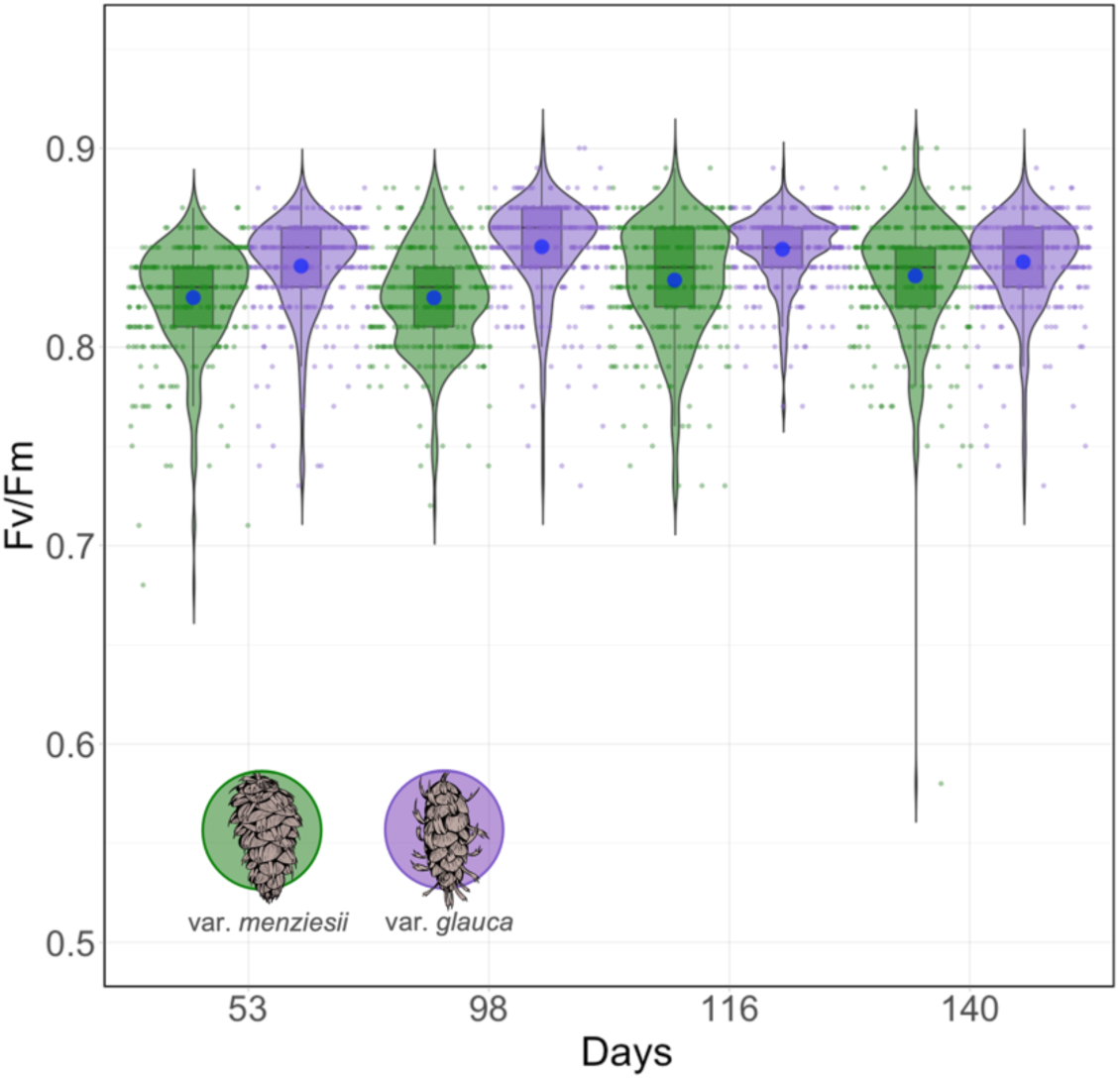
Four individual F_v_/F_m_ measurements of all var. *menziesii* (green) and var. *glauca* (purple) seedlings (points) in the control treatment. Blue dots depict the F_v_/F_m_ average for the control treatment within each variety. These measurements were not corrected for spatial autocorrelation. See Table S3 for statistical test results.

**Fig. S9.**
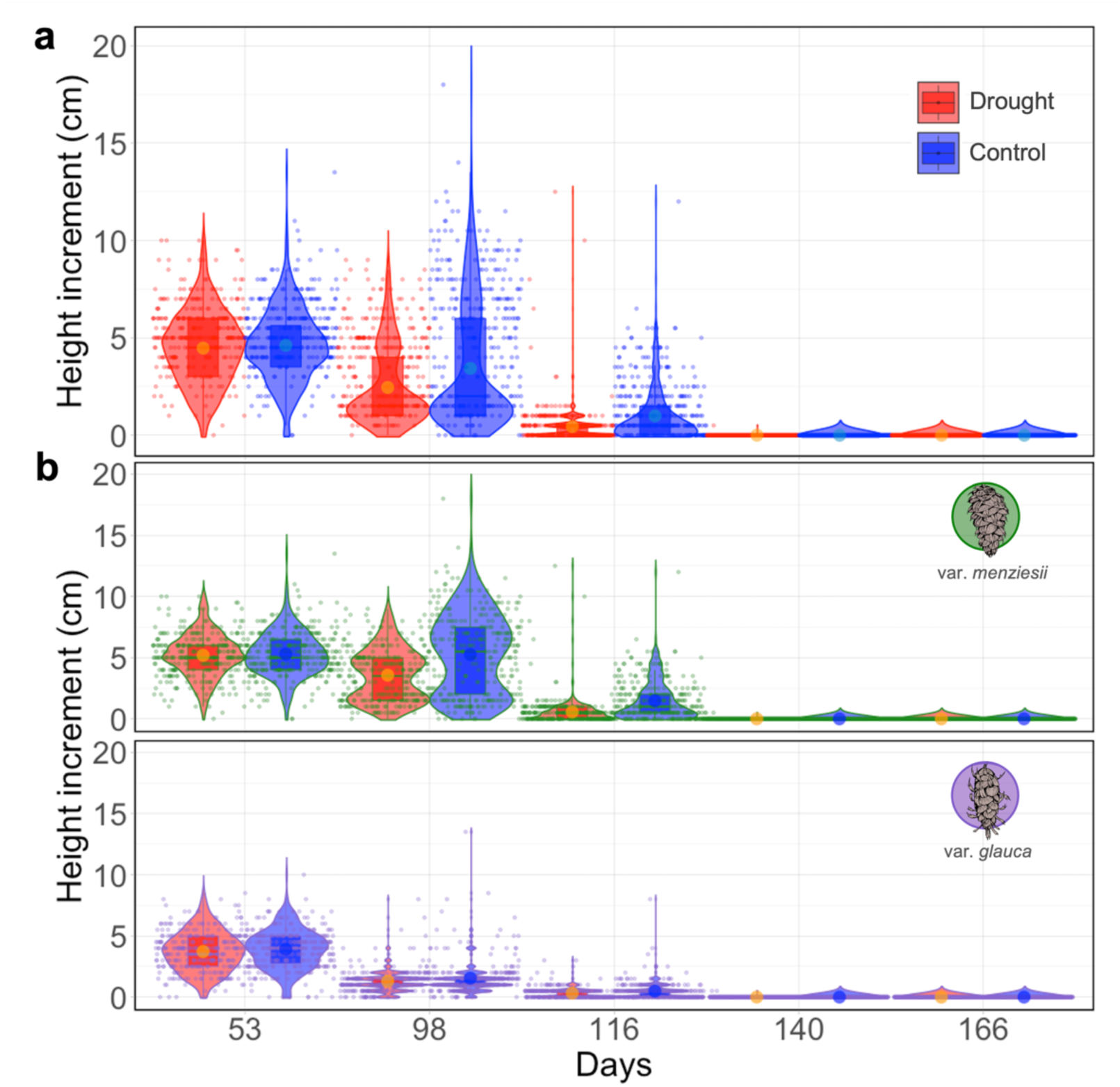
Effect of treatment (control vs. drought) on individual height increment measurements. Species-wide (**a**) and within varieties (**b**) level testing. Orange and blue dots depict the height increment averages for the drought and control treatments respectively. See Table S2 for statistical test results.

**Fig. S10.**
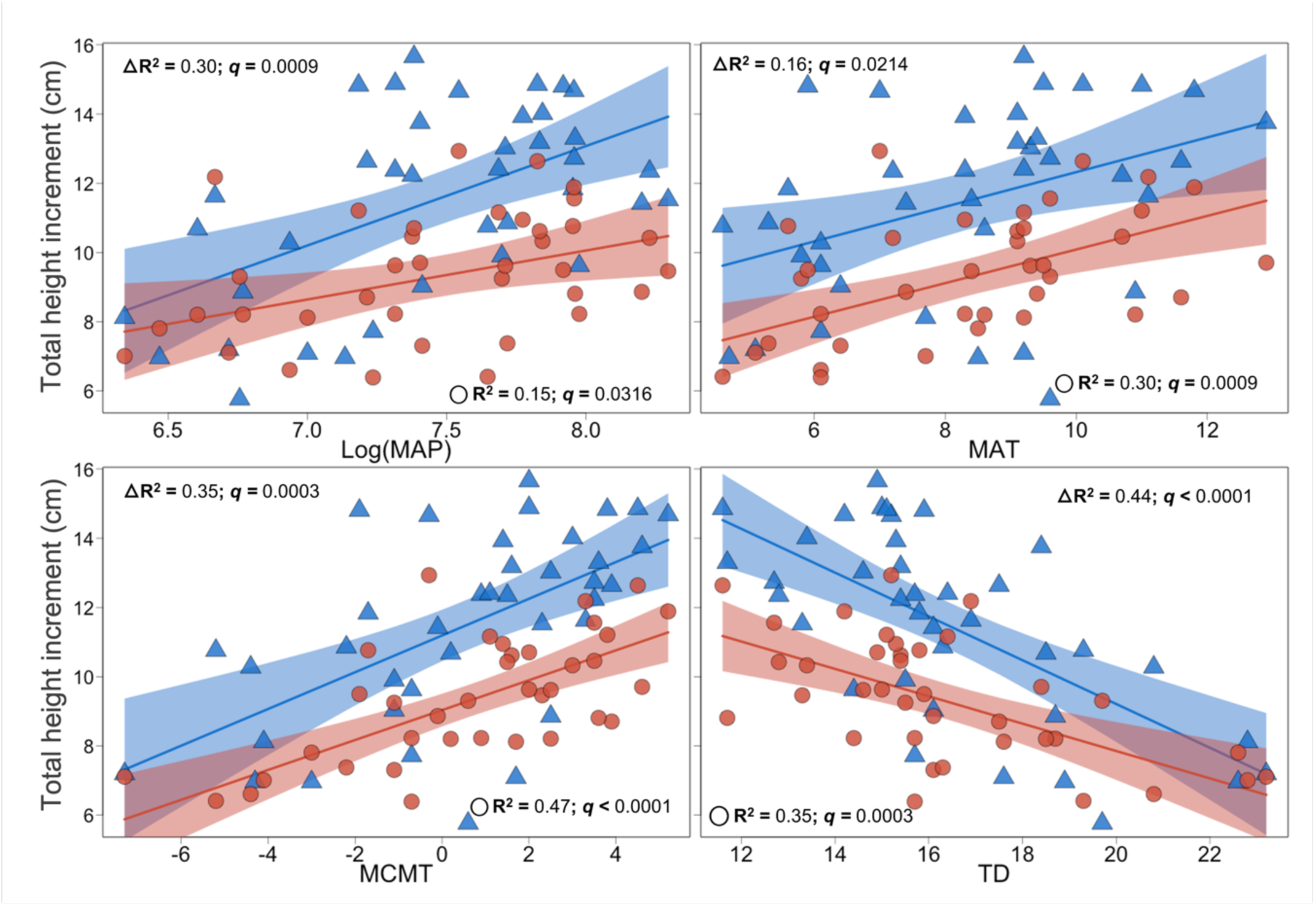
Relationships between total height increment (THI (cm)) BLUEs in drought (red circles) and control (blue triangles) treatments and the var. *menziesii* climate spaces (within variety level). Confidence intervals (95%) of significant relationships (*q-value* < 0.05) are depicted as ribbons. *q-value* is the same as the false-discovery-rate-adjusted *p-value* to account for multiple comparisons. Climatic variables shown: mean annual precipitation (log (MAP (mm))), mean annual temperature (MAT (°C)), mean coldest month temperature (MCMT (°C)), continentality (TD (°C)). See table S6 for results with the other selected environmental variables.

**Fig. S11.**
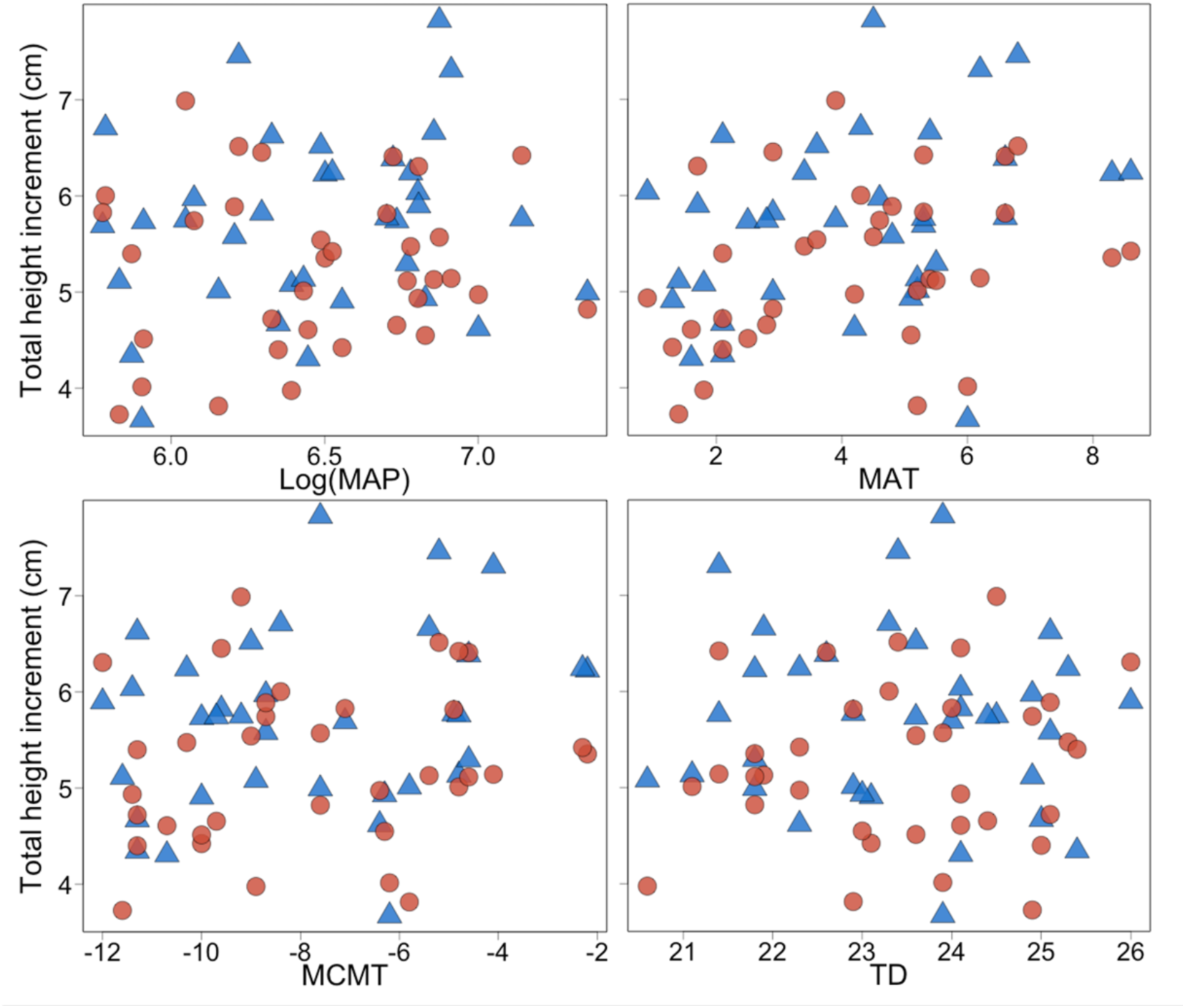
Relationships between total height increment (THI (cm)) BLUEs in drought (red circles) and control (blue triangles) treatments and the var. *glauca* climate spaces (within variety level). No significant relationship was found. Climatic variables shown: mean annual precipitation (log (MAP (mm))), mean annual temperature (MAT (°C)), mean coldest month temperature (MCMT (°C)), continentality (TD (°C)). See table S6 for results with the other selected environmental variables.

**Fig. S12.**
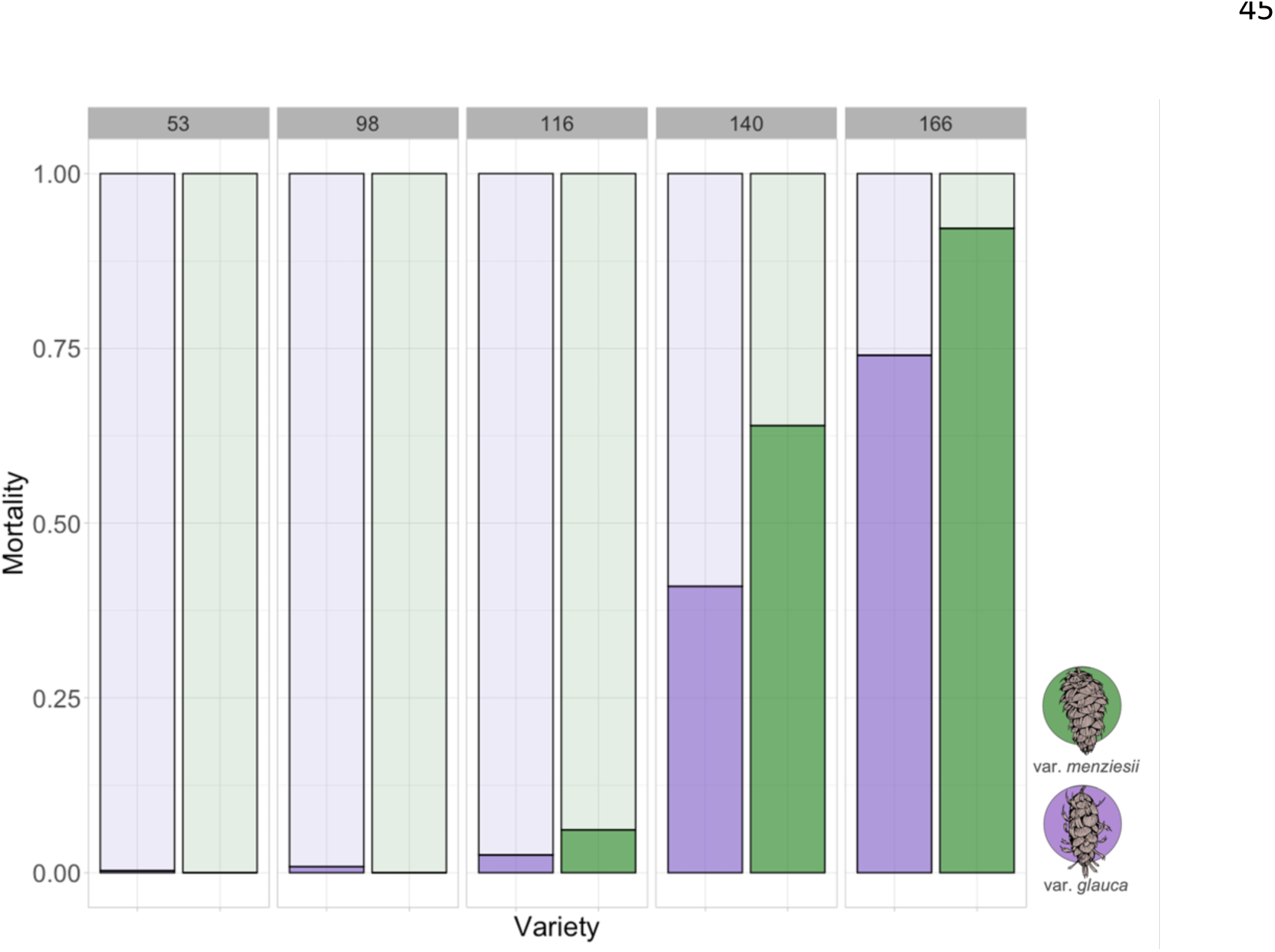
Proportion of mortality within varieties over time (53 to 166 days) in the drought treatment.

